# Modelling palaeoecological time series using generalized additive models

**DOI:** 10.1101/322248

**Authors:** Gavin L. Simpson

**Affiliations:** Institute of Environmental Change and Society, University of Regina, Regina, Saskatchewan, Canada, S4S 0A2

**Keywords:** time series, generalized additive model, simultaneous interval, spline, environmental change

## Abstract

In the absence of annual laminations, time series generated from lake sediments or other similar stratigraphic sequences are irregularly spaced in time, which complicates formal analysis using classical statistical time series models. In lieu, statistical analyses of trends in palaeoenvironmental time series, if done at all, have typically used simpler linear regressions or (non-) parametric correlations with little regard for the violation of assumptions that almost surely occurs due to temporal dependencies in the data or that correlations do not provide estimates of the magnitude of change, just whether or not there is a linear or monotonic trend. Alternative approaches have used Loess-estimated trends to justify data interpretations or test hypotheses as to the causal factors without considering the inherent subjectivity of the choice of parameters used to achieve the Loess fit (e.g. span width, degree of polynomial). Generalized additive models (GAMs) are statistical models that can be used to estimate trends as smooth functions of time. Unlike Loess, GAMs use automatic smoothness selection methods to objectively determine the complexity of the fitted trend, and as formal statistical models, GAMs, allow for potentially complex, non-linear trends, a proper accounting of model uncertainty, and the identification of periods of significant temporal change. Here, I present a consistent and modern approach to the estimation of trends in palaeoenvironmental time series using GAMs, illustrating features of the methodology with two example time series of contrasting complexity; a 150-year bulk organic matter *δ*^15^N time series from Small Water, UK, and a 3000-year alkenone record from Braya-S*ϕ*, Greenland. I discuss the underlying mechanics of GAMs that allow them to learn the shape of the trend from the data themselves and how simultaneous confidence intervals and the first derivatives of the trend are used to properly account for model uncertainty and identify periods of change. It is hoped that by using GAMs greater attention is paid to the statistical estimation of trends in palaeoenvironmental time series leading to more a robust and reproducible palaeoscience.

## 1 Introduction

Palaeoecology and palaeolimnology have moved away from being descriptive disciplines, rapidly adopting new statistical developments in the 1990s and beyond (Smol et al., 2012). Less development has been observed in the area of trend estimation in palaeoenvironmental time series. The vast majority of data produced by palaeoecologists and palaeolimnologists is in the form of time-ordered observations on one or more proxies or biological taxa (Birks, 2012b; Smol, 2008; Smol et al., 2012). Typically these data are arranged irregularly in time; in the absence of annual laminae or varves, the sediment core is sectioned at regular depth intervals (Glew et al., 2001), which, owing to variation in accumulation rates over time and compaction by overlying sediments, results in an uneven sampling in time. An under-appreciated feature of such sampling is that younger sediments often have larger variance than older sediments; each section of core represents fewer lake years in newer samples, relative to older samples. This variable averaging acts as a time-varying low-pass (high-cut) filter of the annual depositional signal.

Irregular intervals between samples means that the time-series analysis methods of autoregressive or moving average processes, in the form of autoregressive integrated moving average (ARIMA) models, are practically impossible to apply because software typically requires even spacing of observations in time. Dutilleul et al. (2012) and Birks (2012a), eschewing the term *time series*, prefer to call such data *temporal series* on account of the irregular spacing of samples, a distinction that I find unnecessary, however.

Where statistical approaches have been applied to trend estimation in palaeoenvironmental time series, a commonly-used method is Loess (Birks, 1998, 2012a; Cleveland, 1979; Juggins and Telford, 2012). Loess, locally weighted scatterplot smoother, as it’s name suggests, was developed to smooth x-y scatterplot data (Cleveland, 1979). The method fits a smooth line through data by fitting weighted least squares (WLS) models to observations within a user-specified window of the focal point, whose width is typically expressed as a proportion *α*. of the *n* data points. Weights are determined by how close (in the x-axis only) an observation in the window is to the focal point giving greatest weight given to points closest to the focal point. The interim Loess-smoothed value for the focal point is the predicted value from the weighted regression at the focal point. The interim values are updated using weights based on how far in the y-axis direction the interim smoothed value lies from the observed value plus the x-axis distance weights; this has the effect of down-weighting outlier observations. The final Loess is obtained by joining the smoothed values. The user has to choose how large a window to use, whether to fit degree 1 (linear) or degree 2 (quadratic) polynomials in the WLS model, and how to weight points in the x-axis. When used in an exploratory mode, the user has considerable freedom to choose the detail of the Loess fit; the window width, for example, can be infinitely tweaked to give as close a fit to the data, as assessed by eye, as is desired. Using cross-validation (CV) to choose *α* or the degree of polynomial in the WLS model is complicated for a number of reasons, not least because the CV scheme used must involve the time ordering of the data (e.g. Bergmeir et al., 2018). This subjectivity is problematic however once we wish to move beyond exploratory analysis and statistically identify trends to test hypotheses involving those trend estimates.

Running means or other types of filter (Juggins and Telford, 2012) have also been used extensively to smooth palaeoenvironmental time series, but, as with Loess, their behaviour depends on a number of factors, including the filter width. Furthermore, the width of the filter causes boundary issues; with a centred filter, of width five, the filtered time series would be two data points shorter at both ends of the series because the filter values are not defined for the first and last two observations of the original series as these extra time points were not observed. Considerable research effort has been expended to identify ways to pad the original time series at one or both ends to maintain the original length in the filtered series, without introducing bias due to the padding (e.g. Mann, 2004, 2008; Mills, 2006, 2007, 2010).

These are not the only methods that have been used to estimated trends in stratigraphic series. Another common approach involves fitting a simple linear trend using ordinary least squares regression and use the resulting *t* statistic as evidence against the null hypothesis of no trend despite the statistical assumptions being almost surely violated due to dependence among observations. The Pearson correlation coefficient, *r*, is also often used to detect trends in palaeo time series (Birks, 2012a), despite the fact that *r* provides no information as to the magnitude of the estimated trend, and the same temporal autocorrelation problem that dogs ordinary least squares similarly plagues significance testing for *r* (Tian et al., 2011). Additionally, both the simple least squares trend line and *r* are tests for *linear* trends only, and yet we typically face data sets with potentially far more complex trends than can be identified by these methods. Instead, non-parametric rank correlation coefficients have been used (Birks, 2012a; Gautheir, 2001), and whilst these do allow for the detection of non-linear trends, trends are restricted to be monotonic, no magnitude of the trend is provided, and the theory underlying significance testing of Spearman’s *ρ* and Kendall’s *τ* assumes independent observations.

Here, I describe generalized additive models (GAMs; Hastie and Tibshirani, 1986,1990; Rup-pert et al., 2003; Wood, 2017; Yee and Mitchell, 1991) for trend estimation. GAMs, like simple linear regression, are a regression-based method for estimating trends, yet they are also, superficially at least, similar to Loess. GAMs and Loess estimate smooth, non-linear trends in time series and both can handle the irregular spacing of samples in time, yet GAMs do not suffer from the subjectivity that plagues Loess as a method of formal statistical inference.

In the subsequent sections, I present an introduction to GAMs and discuss the issue of uncertainty in model-estimated trends, the topic of posterior simulation from a regression model and how to identify periods of significant environmental change using the first derivative of the estimated trend. The main steps in the analysis of palaeoenvironmental time series using GAMs are illustrated in Figure 1. Two non-standard types of spline — adaptive smoothers and Gaussian process splines — that are especially applicable to GAMs in the palaeoenvironmental setting are subsequently described, followed by an assessment of the the impact of age-model uncertainty on trend estimation via GAMs. Finally, I briefly discuss the application of GAM trend analysis to multivariate species abundance and compositional data.

**Figure 1:**
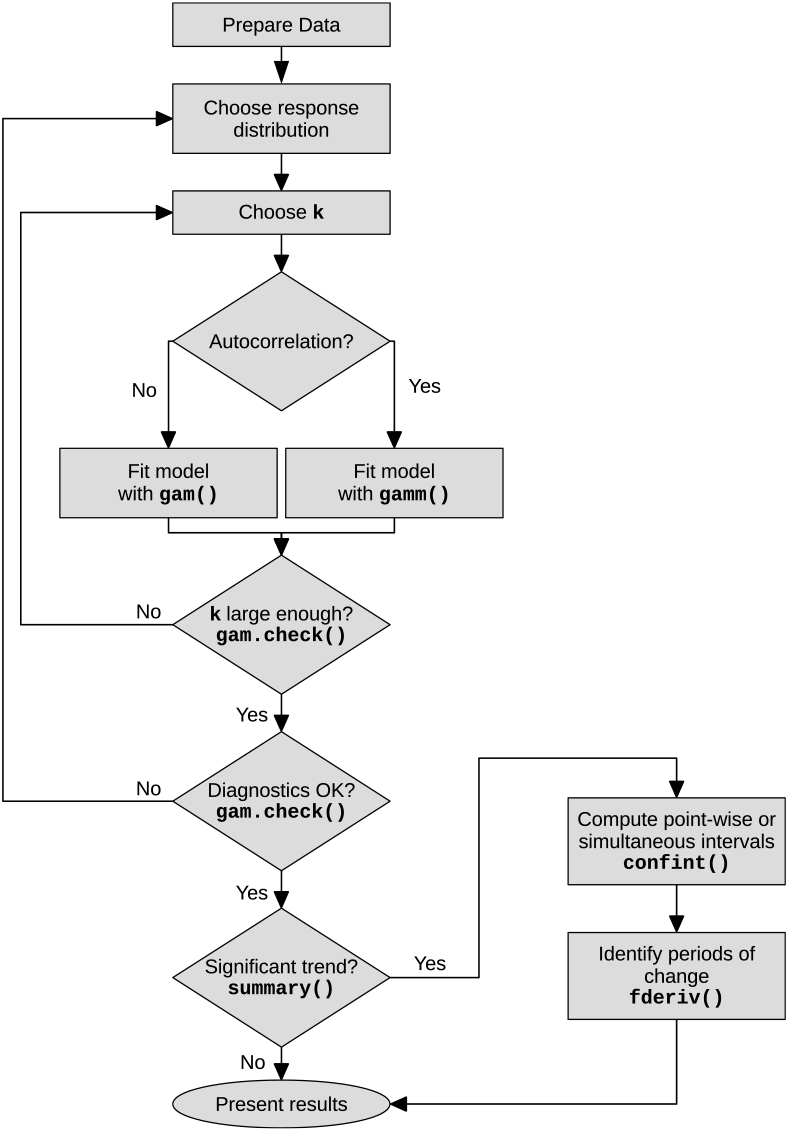
Flowchart showing the main steps in the analysis of time series using generalized additive models. The main R functions associated with each step or decision are shown in bold.

### 1.1 Example time series

To illustrate trend estimation in palaeoenvironmental data using GAMs, I use two proxy time series; a 150-year bulk organic matter *δ*^15^N record from Small Water, and a 3000-year alkenone record from Braya-S*ϕ*. Between them, the two examples, combine many of the features of interest to palaeoecologists that motivate the use of GAMs; non-linear trends and the question of when changes in the measured proxy occurred. The example analyses were all performed using the *mgcv* package (version 1.8.24; Wood, 2017) and R (version 3.4.4; R Core Team, 2018), and the supplementary material contains a fully annotated document showing the R code used to replicate all the analyses described in the remainder of the paper.

#### 1.1.1 *δ*^15^N time series from Small Water

Figure 2a shows 48 nitrogen stable isotope measurements on the bulk organic matter of a sediment core collected from Small Water, a small corrie lake located in the English Lake District, UK. The data were collected to investigate disturbance of nitrogen (N) cycling in remote, oligotrophic lakes by N deposited from the atmosphere (Simpson, unpublished data). The data are shown on a ^210^Pb time scale. Questions that might be asked about this series are; what is the trend in *δ*^15^N?, when do we first see evidence for a change in *δ*^15^N?, and is the reversal in *δ*^15^N values in the uppermost section of the core a real change?

**Figure 2:**
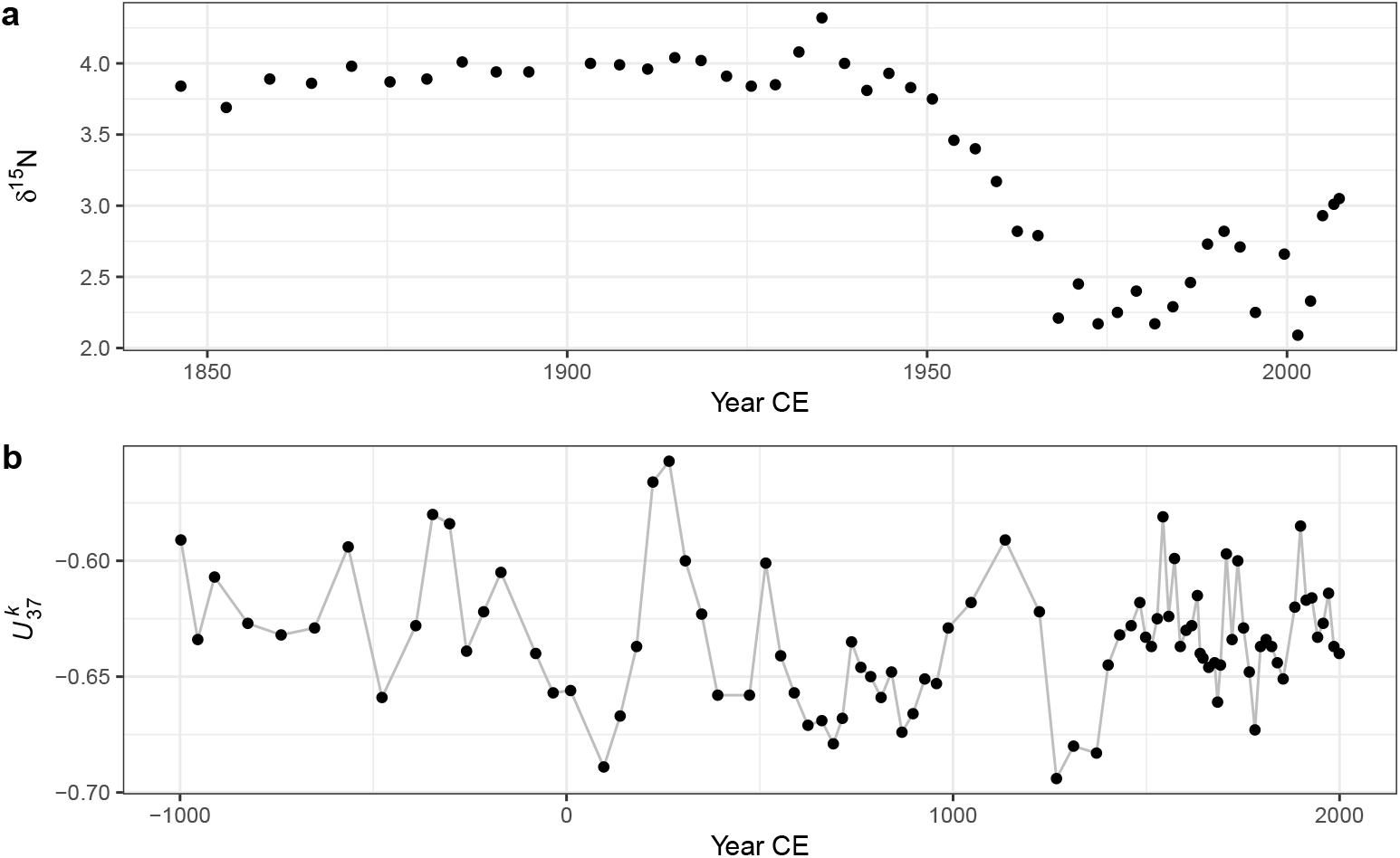
Example time series; a) Small Water bulk organic matter *δ*^15^N time series on a ^210^Pb time scale, and b) Braya-S*ϕ* 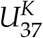 time series on a calibrated ^14^C time scale. The observations 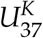 time series have been joined by lines purely as a visual aid to highlight potential trends.

#### 1.1.2 Braya-S*ϕ* alkenone time series

The second example time series is a 3,000 year record of alkenone unsaturation, 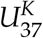, from Braya-S*ϕ*, a meromictic lake in West Greenland (D’Andrea et al., 2011). Alkenones are long-chained unsaturated organic compounds that are produced by a small number of planktonic organisms known as haptophytes. The 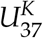 unsaturation index (Brassell, 1993) is

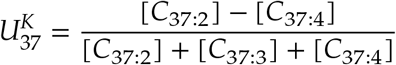

where [*C*_37:*x*_] is the concentration of the alkenone with 37 carbon atoms and *x* double carbon bonds. The relative abundance of these alkenones is known to vary with changes in water temperature (Brassell, 1993; Chu et al., 2005; Toney et al., 2010; Zink et al., 2001), and as a result 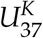 is used as a proxy for lake- and sea-surface temperatures. For further details on the Braya-S*ϕ* 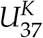 record and age model see D’Andrea et al. (2011). Here I use the 3,000 year 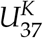 record from the PAGES 2K database (PAGES 2K Consortium, 2013). The data are presented in Figure 2b.

## 2 Regression models for palaeoenvironmental time series

A linear model for a trend in a series of *T* observations *y_t_* at observation times *x_t_* with *t* = 1,2,…, *T* is

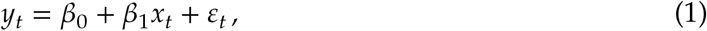

where *β*_0_ is a constant term, the model *intercept*, representing the expected value of *y_t_* where *x_t_* is 0. *β*_1_ is the *slope* of the best fit line through the data; it measures the rate of change in *y* for a unit increase in *x*. The unknowns, the *β_j_*, are commonly estimated using least squares by minimising the sum of squared errors, 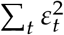. If we want to ask if the estimated trend *β*_1_ is statistically significant we must make further assumptions about the data (conditional upon the fitted model) or the model errors (residuals); 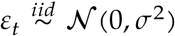. This notation indicates that the residuals *ϵ_t_* are *independent* and *identically distributed* Gaussian random variables with mean equal to 0 and constant variance *σ*^2^. In the time series setting, the assumption of independence of model residuals is often violated.

The linear model described above is quite restrictive in terms of the types of trend it can fit; essentially linear increasing or decreasing trends, or, trivially, a null trend of no change. This model can be extended to allow for non-linear trends by making *y_t_* depend on polynomials of *x_t_*, for example

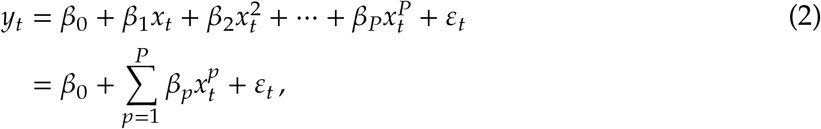

where polynomials of *x_t_* up to order *P* are used. This model allows for more complex trends but it remains a fully parametric model and suffers from several problems, especially the behaviour of the fitted trend at the start and end of the observed series.

Linear models using a range of polynomials fitted to the Small Water data set are shown in Figure 3. The low-order models (*P* ∈ {1, 3}) result in very poor fit to the data. The model with *P* = 5 does a reasonable job of capturing the gross pattern in the time series, but fails to adapt quickly enough to the decrease in *δ*^15^N that begins ~1940 CE, and the estimated trend is quite biased as a result. The *P* = 10th-order polynomial model is well able to capture this period of rapid change, but it does so at the expense of increased complexity in the estimated trend prior to ~1940. Additionally, this model (*P* = 10) has undesirable behaviour at the ends of the series, significantly overfitting the data, a commonly observed problem in polynomial models such as these (Epperson, 1987; Runge, 1901). Finally, the choice of what order of polynomial to fit is an additional choice left for the analyst to specify; choosing the optimal *P* is not a trivial task when the data are a time series and residual autocorrelation is likely present.

**Figure 3:**
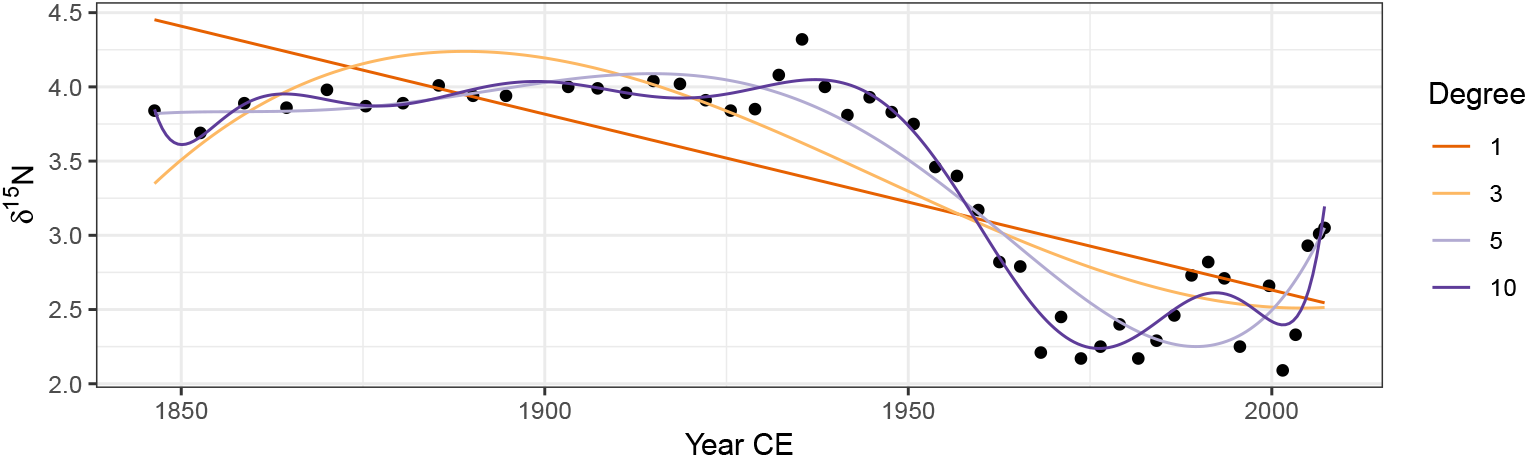
Linear models with various orders of polynomial of the covariate Year fitted using ordinary least squares to the *δ*^15^N time series from Small Water. The degree of polynomial is indicated, with the degree 1 line equal to a simple linear regression model.

Can we do better than these polynomial fits? In the remainder, I hope to demonstrate that the answer to that question is emphatically “yes”! Below I describe a coherent and consistent approach to modelling palaeoenvironmental time series using generalized additive models that builds upon the linear regression framework.

## 3 Generalized additive models

The GAM version of the linear model (1) is

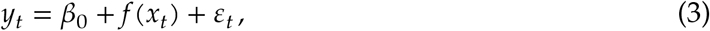

where the linear effect of time (the *β*_1_*x_t_* part) has been replaced by a smooth function of time, *f*(*x_t_*). The immediate advantage of the GAM is that we are no longer restricted to the shapes of trends that can be fitted via global polynomial functions such as (2). Instead, the shape of the fitted trend will be estimated from the data itself.

The linear model is a special case of a broader class, known as the generalized linear model (GLM; McCullagh and Nelder, 1989). The GLM provides a common framework for modelling a wide range of types of data, such as count, proportions, or binary (presence/absence) data, that are not conditionally distributed Gaussian. GLMs are, like the linear model, parametric in nature; the types of trends that we can fit using a GLM are the linear or polynomial models. GAMs extend the GLM by relaxing this parametric assumption; in a GAM some, or all, of the parametric terms, the *β_p_*, are replace by smooth functions *f_j_* of the covariates *x_j_*. For completeness then, we can write (3) as a GLM/GAM

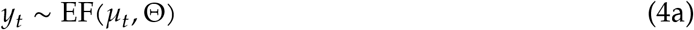

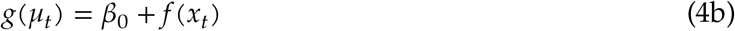

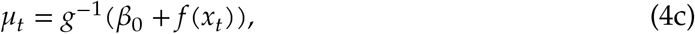

where *μ_t_* is the expected value (e.g. the mean count or the probability of occurrence) of the random variable 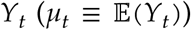 of which we have observations *y_t_*. *g* is the link function, an invertible, monotonic function, such as the natural logarithm, and *g*^−1^ is its inverse. The link function maps values from the response scale on to the scale of the linear predictor, whilst the inverse of the link function provides the reverse mapping. For example, count data are strictly non-negative integer values and are commonly modelled as a Poisson GLM/GAM using the natural log link function. On the log scale, the response can take any real value between −∞ and +∞, and it is on this scale that model fitting actually occurs (i.e. using equation (4b)). However we need to map these unbounded values back on to the non-negative response scale. The inverse of the log link function, the exponential function, achieves this and maps values to the interval 0-∞ (equation (4c)).

In (4a), we further assume that the observations are drawn from a member of the exponential family of distributions — such as the Poisson for count data, the binomial for presence/absence or counts from a total — with expected value *μ_t_* and possibly some additional parameters Θ (*y_t_* ~ EF (*μ_t_*, Θ)). Additionally, many software implementations of the above model also allow for distributions that are not within the exponential family but which can be fitted using an algorithm superficially similar to the one used to fit GAMs to members of the exponential family (e.g. Wood et al., 2016). Common examples of such extended families include the negative binomial distribution (for overdispersed counts) and the beta distribution (for true proportions or other interval-bounded data).

### 3.1 Basis functions

It is clear from plots of the data (Figure 2) that we require the fitted trends for the Small Water *δ*^15^N and Braya-S*ϕ* 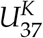 time series to be non-linear functions, but it is less clear how to specify the actual shape require. Ideally, we’d like to learn the shape of the trend from the data themselves. We will refer to these non-linear functions as *smooth functions*, or just *smooths* for short, and we will denote a smooth using *f*(*x_t_*). Further, we would like to represent the smooths in a way that (4) is represented parametrically so that it can be estimate within the well-studied GLM framework. This is achieved by representing the smooth using a *basis*. A basis is a set of functions that collectively span a space of smooths that, we hope, contains the true *f*(*x_t_*) or a close approximation to it. The functions in the basis are known as *basis functions*, and arise from a *basis expansion* of a covariate. Writing *b_j_*(*x_t_*) as the *j*th basis function of *x_t_*, the smooth *f*(*x_t_*) can be represented as a weighted sum of basis functions

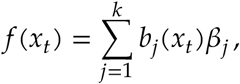

where *β_j_* is the weight applied to the *j*th basis function.

The polynomial model is an example of a statistical model that uses a basis expansion. For the cubic polynomial (*P* = 3) fit shown in Figure 3 there are in fact 4 basis functions: 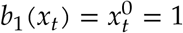, *b*_2_(*x_t_*) = *x_t_*, 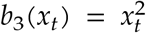, and 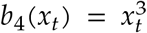. Note that *b*_1_(*x_t_*) is constant and is linked to the model intercept, *β*_0_, in the linear model (2), and further, that the basis function weights are the estimated coefficients in the model, the *β_j_*.

As we have already seen, polynomial basis expansions do not necessarily lead to well-fitting models unless the true function *f* is itself a polynomial. One of the primary criticisms is that polynomial basis functions are global (Magee, 1998); the value of *f* at time point *x_t_* affects the value of *f* at time point *x*_*t*+*s*_ even if the two time points are at opposite ends of the series. There are many other bases we could use; here I discuss one such set of bases, that of splines.

There are a bewildering array of different types of spline. In the models discussed below we will largely restrict ourselves to cubic regression splines (CRS) and thin plate regression splines (TPRS). In addition, I also discuss two special types of spline basis, an adaptive spline basis and a Gaussian process spline basis.

A cubic spline is a smooth curve comprised of sections of cubic polynomials, where the sections are joined together at some specified locations — known as *knots* — in such a way that at the joins, the two sections of cubic polynomial that meet have the same value as well as the same first and second derivative. These properties mean that the sections join smoothly and differentiably at the knots (Wood, 2017,5.3.1).

The CRS can be parameterized in a number of different ways. One requires a knot at each unique data value in *x_t_*, which is computationally inefficient. Another way of specifying a CRS basis is to parameterize in terms of the value of the spline at the knots. Typically in this parametrization there are many fewer knots than unique data, with the knots distributed evenly over the range of *x_t_* or at the quantiles of *x_t_*. Placing knots at the quantiles of *x_t_* has the effect of placing a greater number of knots where the data is most dense.

A CRS basis expansion comprised of 7 basis functions for the time covariate in the Small Water series, is shown in Figure 4a. The tick marks on the x-axis show the locations of the knots, which are located at the ends of the series and evenly in between. Notice that in this particular parametrization, the *j*th basis function takes a value of 1 at the *j*th knot and at all other knots a value of 0.

**Figure 4:**
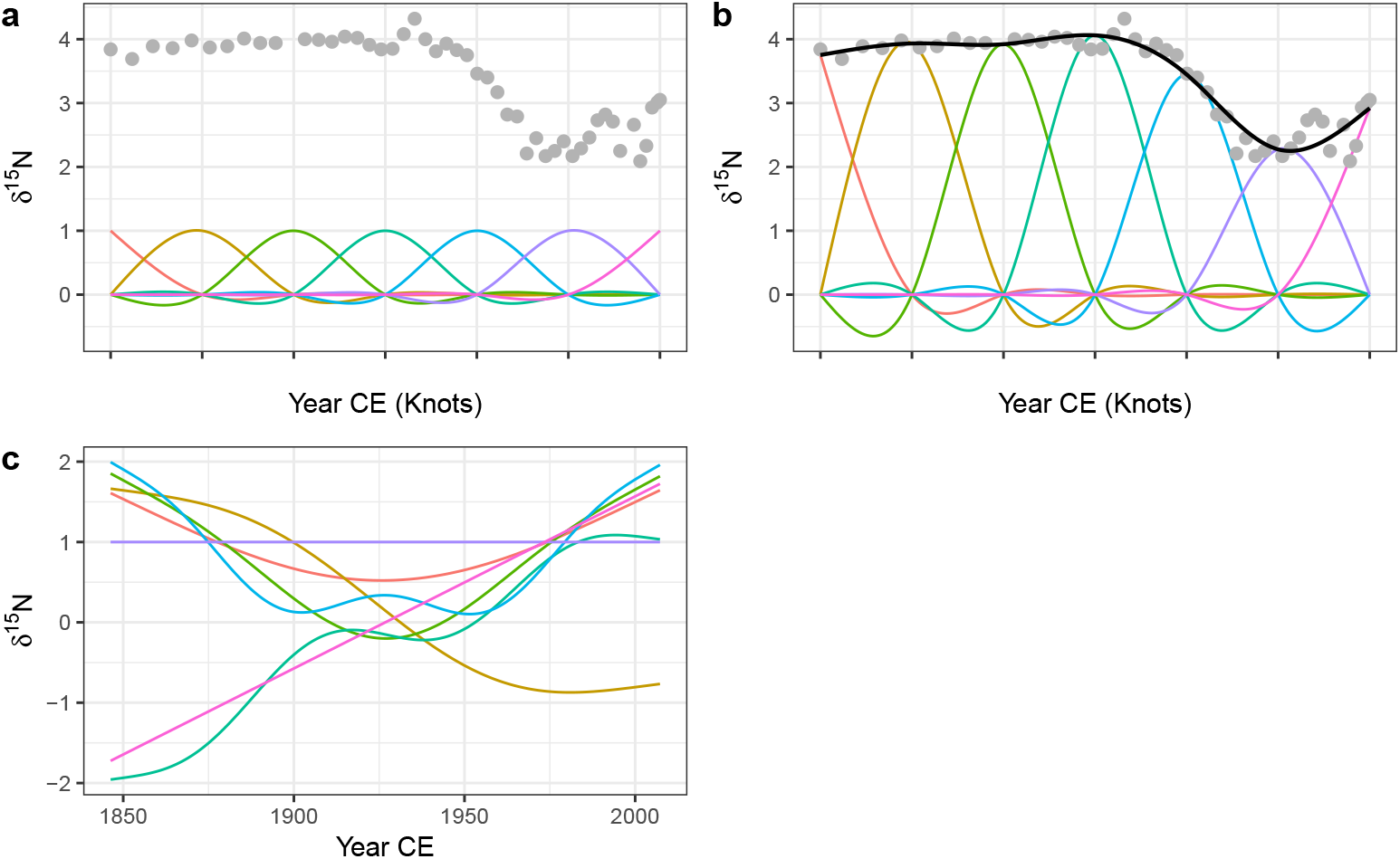
Basis functions for the time covariate and the Small Water *β*^15^N time series. A rank (size) 7 cubic regression spline (CRS) basis expansion is show in a), with knots, indicated by tick marks on the x-axis, spread evenly through the rang of the data. b) shows the same CRS basis functions weighted by the estimated coefficients *β_j_*, plus the resulting GAM trend line (black line drawn through the data). The grey points in both panels are the observed *β*^15^N values. c) A rank 7 thin plate regression spline basis for the same data.

To estimate a model using this basis expansion each basis function forms a column in the model matrix **X** and the weights *β_j_* can be found using least squares regression (assuming a Gaussian response). Note that in order to estimate a coefficient for each basis function the model has to be fitted without an intercept term. In practice we would include an intercept term in the model and therefore the basis functions are modified via an identifiability constraint (Wood, 2017). This has the effect of making the basis orthogonal to the intercept but results in more complicated basis functions than those shown in in Figure 4a.

Having estimated the weight for each basis function, the *j*th basis function *b_j_* is scaled (weighted) by its coefficient *β_j_*. The scaled CRS basis functions for the Small Water time series are shown in Figure 4b. The solid line passing through the data points is formed by summing up the values of the seven scaled basis functions (*b_j_*(*x_t_*)*β_j_*) at any value of *x_t_* (time).

Cubic regression splines, as well as many other types of spline, require the analyst to choose the number and location of the knots that parametrise the basis. Thin plate regression splines (TPRS) remove this element of subjectivity when fitting GAMs. Thin plate splines were introduced by Duchon (1977) and, as well as solving the knot selection problem, have several additional attractive properties in terms of optimality and their ability to estimate a smooth function of two or more variables, leading to smooth interactions between covariates. However, thin plate splines have one key disadvantage over CRS; thin plate splines have as many unknown parameters as there are unique combinations of covariate values in a data set (Wood, 2017, 5.5.1). It is unlikely that any real data problem would involve functions of such complexity that they require as many basis functions as data. It is much more likely that the true functions that we attempt to estimate are far simpler than the set of functions representable by 1 basis function per unique data value. From a practical point of view, it is also highly inefficient to carry around all these basis functions whilst model fitting, and the available computational resources would become quickly exhausted for large time series with many observations.

To address this issue low rank thin plate regression splines (TPRS) have been suggested which truncate the space of the thin plate spline basis to some lower number of basis functions whilst preserving much of the advantage of the original basis as an optimally-fitting spline (Wood, 2003). A rank 7 TPRS basis (i.e. one containing 7 basis functions) is shown in Figure 4c for the Small Water time series. The truncation is achieved by performing an eigen-decomposition of the basis functions and retaining the eigenvectors associated with the *k* largest eigenvalues. This is similar to the way principal components analysis decomposes a data set into axes of variation (eigenvectors) in decreasing order of variance explained. The truncated basis can preserve much of the space of functions spanned by the original basis but at the cost of using far fewer basis functions (Wood, 2003, 2017, 5.5.1). Note the horizontal TPRS basis function (at *δ*^15^N = 1) in Figure 4c; this basis function is confounded with the intercept term and, after the application of identifiability constraints, ends up being removed from the set of basis functions used to fit the model.

The truncation suggested by Wood (2003) is not without cost; the eigen-decomposition and related steps can be relatively costly for large data sets. For data sets of similar size to the two examples used here, the additional computational effort required to set up the TPRS basis over the CRS basis will not be noticeable. For highly resolved series containing more than ~1000 observations the truncation may be costly computationally. In such instances, little is lost by moving to the CRS basis with the same number of knots as the rank of the desired TPRS, with the benefit of considerably reduced set up time for the basis.

To fit a GAM using either of the two regression spline bases described above, the analyst is generally only required to the specify the size (rank) of the basis expansion required to represent or closely approximate the true function *f*. With practice and some knowledge of the system from which the observations arise, it can be relatively easy to put an upper limit on the expected complexity of the true trend in the data. Additionally, the number of available data points places a constraint on the upper limit of the size of basis expansion that can be used.

In practice, the size of the basis is an upper limit on the expected complexity of the trend, and a simple test can be used to check if the basis used was sufficiently large (Pya and Wood, 2016). This test is available via the `gam.check()` function in *mgcv* for example, which essentially looks at whether there is any additional nonlinearity or structure in the residuals that can be explained by a further smooth of *x_t_*. Should a smooth term in the fitted model fail this test the model can be refitted using a larger basis expansion, say by doubling the value of *k* (the rank) used to fit the original. Note also that a smooth might fail this test whilst using fewer effective degrees of freedom than the maximum possible for the dimension of basis used. This may happen when the true function lies at the upper limit of the set of functions encompassed by the size of basis used. Additionally, a basis of size 2*k* encompasses a richer space of functions of a given complexity than a basis of size *k* (Wood, 2017); increasing the basis dimension used to fit the model may unlock this additional function space resulting in a better fitting model whilst using a similar number of effective degrees of freedom.

### 3.2 Smoothness selection

Having identified low rank regression splines as a useful way to represent *f*, we next need a way to decide how wiggly the fitted trend should be. A backwards elimination approach to sequentially remove knots or basis functions might seem appropriate, however such an approach would likely fail as the resulting sequence of models would not be strictly nested, precluding many forms of statistical comparison (Wood, 2017). Alternatively, we could keep the basis dimension at a fixed size but guard against fitting very complex models through the use of a wiggliness penalty.

The default wiggliness penalty used in GAMs is on the second derivative of the spline, which measures the rate of change of the slope, or the curvature, of the spline at any infinitesimal point in the interval spanned by *x_t_*. The actual penalty used is the integrated squared second derivative of the spline

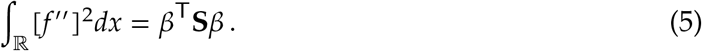

The right hand side of (5) is the penalty in quadratic form. The convenience of the quadratic form is that it is a function of the estimated coefficients of *f*(*x_t_*) where **S** is known as the penalty matrix. Notice that now both the weights for the basis functions and the wiggliness penalty are expressed as functions of the model coefficients.

Now that we have a convenient way to measure wiggliness, it needs to be incorporated into the objective function that will be minimised to fit the GAM. The likelihood of the model given the parameter estimates 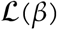 is combined with the penalty to create the penalized likelihood 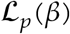:

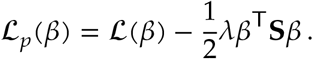

The fraction of a half is there simply to make the penalised likelihood equal the penalised sum of squares in the case of a Gaussian model. *λ* is known as the smoothness parameter and controls the extent to which the penalty contributes to the likelihood of the model. In the extreme case of *λ* = 0 the penalty has no effect and the penalized likelihood equals the likelihood of the model given the parameters. At the other extreme, as *λ* → ∞ the penalty comes to dominate 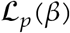 and the wiggliness of *f*(*x_t_*) tends to 0 resulting in an infinitely smooth function. In the case of a second derivative penalty, this is a straight line, and we recover the simple linear trend from (1) when assuming a Gaussian response.

Figure 5 illustrates how the smoothness parameter *λ* controls the degree of wiggliness in the fitted spline. Four models are shown, each fitted with a fixed value of *λ*; 10000, 1, 0.01, and 0.00001. At *λ* = 10000 the model effectively fits a linear model through the data. As the value of *λ* decreases, the fitted spline becomes increasingly wiggly. As *λ* becomes very small, the resulting spline passes through most of the *δ*^15^N observations resulting in a model that is clearly over fitted to the data.

**Figure 5:**
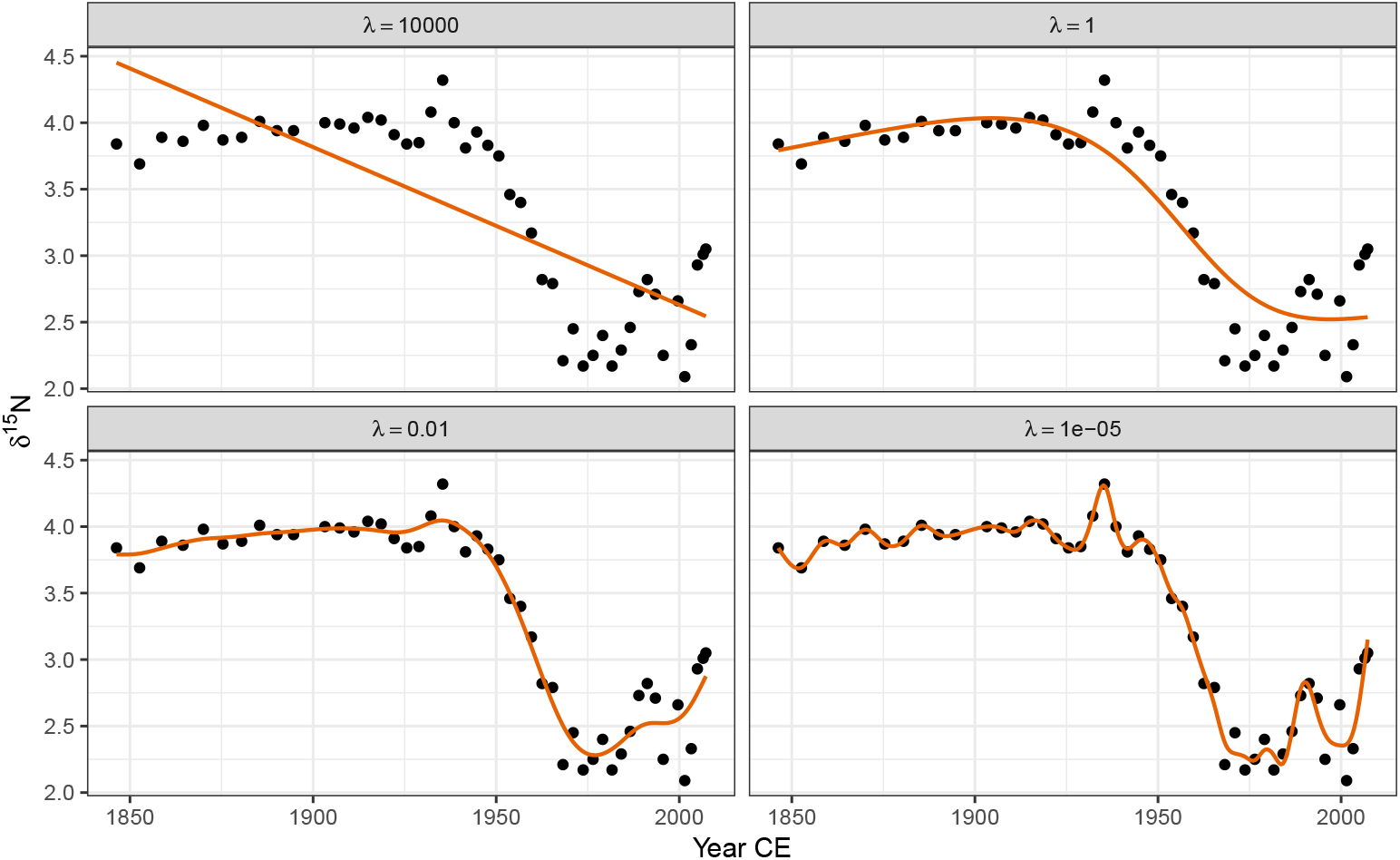
The effect of the smoothness parameter *λ* on the resulting wiggliness of the estimated spline. Large values of *λ* penalize wiggliness strongly, resulting in smooth trends (upper row), while smaller values allow increasingly wiggly trends. The aim of automatic smoothness selection is to find an optimal value of *λ* that balances the fit of the model with model complexity to avoid overfitting.

To fully automate smoothness selection for *f*(*x_t_*) we need to estimate *λ*. There are two main ways that *λ* can be automatically chosen during model fitting. The first way is to choose *λ* such that it minimises the prediction error of the model. This can be achieved by choosing *λ* to minimise Akaike’s information criterion (AIC) or via cross-validation (CV) or generalized cross-validation (GCV; Craven and Wahba, 1978). GCV avoids the computational overhead inherent to CV of having to repeatedly refit the model with one or more observations left out as a test set. Minimising the GCV score will, with a sufficiently large data set, find a model with the minimal prediction error (Wood, 2017). The second approach is to treat the smooth as a random effect, in which *λ* is now a variance parameter to be estimated using maximum likelihood (ML) or restricted maximum likelihood (REML; Wood, 2011; Wood et al., 2016).

Several recent results have shown that GCV, under certain circumstances, has a tendency to under smooth, resulting in fitted splines that are overly wiggly (Reiss and Ogden, 2009). Much better behaviour has been observed for REML and ML smoothness selection, in that order (Wood, 2011). REML is therefore the recommended means of fitting GAMs, though, where models have different fixed effects (covariates) they cannot be compared using REML, and ML selection should be used instead. In the sorts of data examples considered here there is only a single covariate *x_t_* as our models contain a single estimated trend so REML smoothness selection is used throughout unless otherwise stated.

## 4 Fitting GAMs

### 4.1 Small Water

The trend in *δ*^15^N values is clearly non-linear but it would be difficult to suggest a suitable polynomial model that would allow for periods of relatively no change in *δ*^15^N as well as rapid change. Instead, a GAM is ideally suited to modelling such trends; the data suggest a smoothly varying change in *δ*^15^N between 1925 and 1975. It is reasonable to expect some autocorrelation in the model errors about the fitted trend. Therefore I fitted the following GAM to the *δ*^15^N time series.

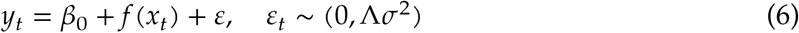

Now the i.i.d. assumption has been relaxed and a correlation matrix, Λ, has been introduced that is used to model autocorrelation in the residuals. The *δ*^15^N values are irregularly spaced in time and a correlation structure that can handle the uneven spacing is needed (Pinheiro and Bates, 2000). A continuous time first-order autoregressive process (CAR(1)) is a reasonable choice; it is the continuous-time equivalent of the first-order autoregressive process (AR(1)) and, simply stated, models the correlation between any two residuals as an exponentially decreasing function of *h*(*ϕ^h^*), where *h* is the amount of separation in time between the residuals (Pinheiro and Bates, 2000). *h* may be a real valued number in the CAR(1), which is how it can accommodate the irregular separation of samples in time. *ϕ* controls how quickly the correlation between any two residuals declines as a function of their separation in time and is an additional parameter that will be estimated during model fitting. The model in (6) was fitted using the `gamm()` function (Wood, 2004) in the *mgcv* package (Wood, 2017) for R (R Core Team, 2017).

The fitted trend is shown in Figure 6a, and well-captures the strong pattern in the data. The trend is statistically significant (estimated degrees of freedom = 7.95; *F* = 47.44, approximate *p* value = ≪ 0.0001). However further analysis of the fitted model is required to answer the other questions posed earlier about the timing of change and whether features in the trend can be distinguished from random noise. I discuss these issues shortly.

**Figure 6:**
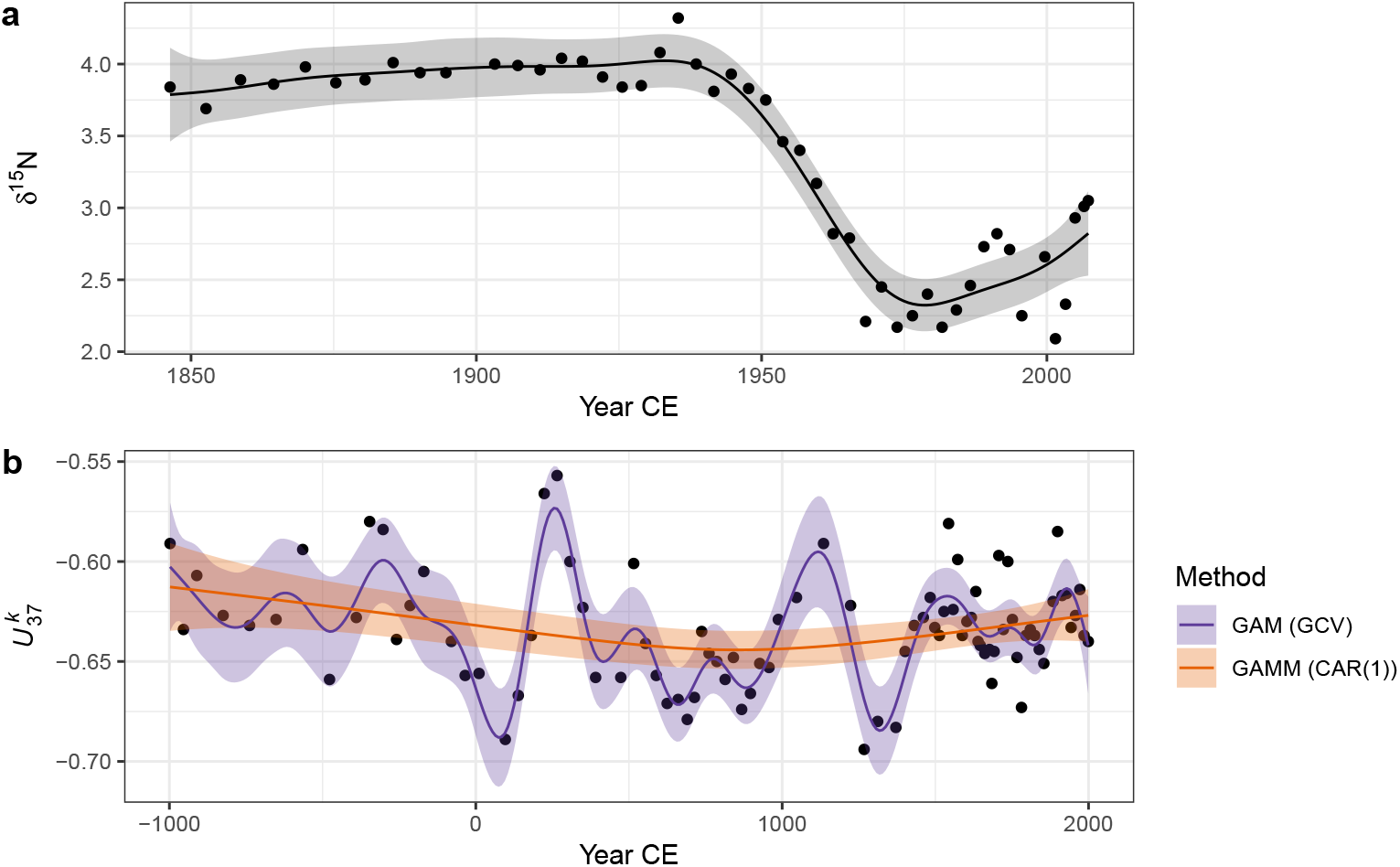
GAM-based trends fitted to the Small Water *δ*^15^N (a) and Braya-S*ϕ* 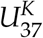 (b) time series. The shaded bands surrounding the estimated trends are approximate 95% across-the-function confidence intervals. For the 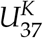 series, two models are shown; the orange fit is the result of a GAM with a continuous-time AR(1) process estimated using REML smoothness selection, while the blue fit is that of a simple GAM with GCV-based smoothness selection. The REML-based fit significantly oversmooths the 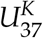 time series.

### 4.2 Braya-S*ϕ*

The 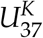 data present a more difficult data analysis challenge than the *δ*^15^N time series because of the much more complex variation present. Fitting the same model as the Small Water example, (6), to the 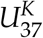 data resulted in the unsatisfactory fit shown as the very smooth line in Figure 6b (labelled GAMM (CAR(1))). Further problems were evident with this model fit — the covariance matrix of the model was non-positive definite, a sure sign of problems with the fitted model. Refitting with a smaller basis dimension (*k* = 20) for the trend term resulted in a model with a positive-definite covariance matrix for the model variance-covariance terms, but the estimated value of of the CAR(1) parameter *ϕ* = 0.2 was exceedingly uncertain (95% confidence interval 0 – 1!)

Fitting this model as a standard GAM with REML smoothness selection resulted in the same fitted trend as the GAM with CAR(1) errors (not shown), whilst using GCV smoothness selection resulted in a much more satisfactory fitted trend. There are two potential problems with the GCV-selected trend: i) GCV is sensitive to the profile of the GCV score and has been shown to under smooth data in situations where the profile is flat around the minimum GCV score, and ii) the model fitted assumes that the observations are independent, an assumption that is certainly violated in the 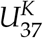 time series.

To investigate the first issue, the GCV and REML scores for an increasing sequence of values of the smoothness parameter (*λ*) were evaluated for the standard GAM (equation (4)) fit to the 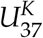 time series. The resulting profiles are shown in Figure 7, with the optimal value of the parameter shown by the vertical line. The GCV score profile suggests that the potential for under smoothing identified by Reiss and Ogden (2009) is unlikely to apply here as there is a well-defined minimum in profile.

**Figure 7:**
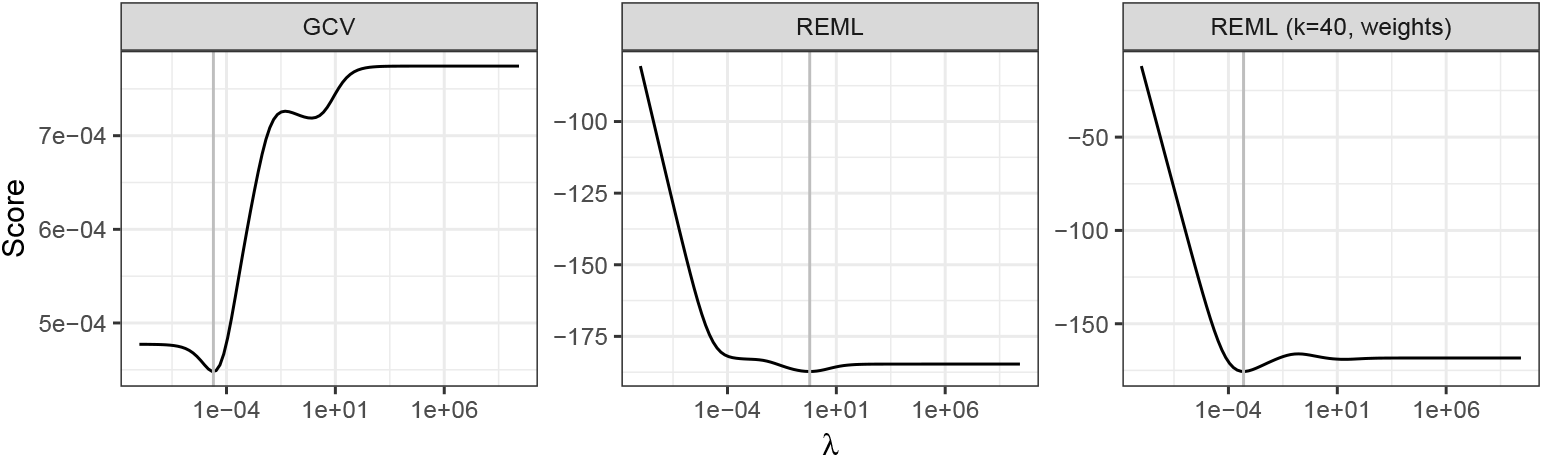
GCV and REML scores as a function of the smoothness parameter *λ*. From left to right, GAMs were estimated using GCV and REML smoothness selection, and REML using a basis dimension of 40 and observational weights to account for heterogeneity in the 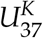 times series. The selected value of *λ* for each model is indicated by the vertical grey line.

To understand the reason why the GAM plus CAR(1) and the simple GAM with REML smoothness selection performed poorly with the 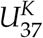 time series we need to delve a little deeper into what is happening when we are fitting these two models.

The primary issue leading to poor fit is that neither model accounts for the different variance (known as (heteroscedasticity) of each observation in the 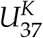 record. This seemingly isn’t a problem for GCV which minimizes prediction error. The sediments in Braya-S*ϕ* are not annually laminated and therefore the core was sliced at regular depth intervals. Owing to compaction of older sediments and variation in accumulation rates over time, each sediment slice represents a different number of “lake years”. We can think of older samples as representing some average of many years of sediment deposition, whilst younger samples are representative of fewer of these lake years. The average of a larger set of numbers is estimated more precisely than the average of a smaller set, all things equal. A direct result of this variable averaging of lake years it that some samples are more precise and therefore have lower variance than other samples and yet the model assumed that the variance was constant across samples.

Accounting for heteroscedasticity within the model is relatively simple via the use of observational weights. The number of lake years represented by each slice is estimated by assigning a date to the top and bottom of each sediment slice. The variance of each observation should be proportional to the inverse of the number of lake years each sample represents. In the `gam()` function used here, weights should be specified as the number of lake years each sample represents. Other software may require the weights to be specified in a different way.

A secondary problem is the size of the basis dimension used for the time variable. The main user selectable option when fitting a GAM in the penalised likelihood framework of Wood (2004) is how many basis functions to use. As described above, the basis should be large enough to contain the true, but unknown, function or a close approximation to it. For GCV selection the basis used contained 29 basis functions, whilst the CAR(1) model with REML smoothness selection would only converge with a basis containing 20 functions. The size of the basis appears to be sufficient for GCV smoothness selection, but following Wood (2011) REML smoothness selection is preferred. Using the test of Pya and Wood (2016), the basis dimension for the models with REML smoothness selection was too small. To proceed therefore, we must drop the CAR(1) term and increase the basis dimension to 39 functions (by setting `k = 40`; I set it larger than expected because the larger basis contains a richer family of functions and the excess complexity is reduced because of the smoothness penalty.)

With the larger basis dimension and accounting for the non-constant variance of the observations via weights, the model fitted using REML is indistinguishable from that obtained using GCV (Figure 6b). The trace of the REML score for this model shows a pronounced minimum at a much smaller value of *λ* than the original REML fit (Figure 7), indicating that a more wiggly trend provides a better fit to the Braya-S*ϕ* time series. This example illustrates that some care and understanding of the underlying principles of GAMs is required to diagnose potential issues with the estimated model. After standard modelling choices (size of basis to use, correct selection of response distribution and link function, etc.) are checked, it often pays to think carefully about the properties of the data and ensure that the assumptions of the model are met. Here, despite increasing the basis size, it was the failure to appreciate the magnitude of the effect of the non-constant variance that lead to the initially poor fit and the problems associated with the estimation of the CAR(1) process. I return to the issue of why the GAM plus CAR(1) model encountered problems during fitting later (see section Residual autocorrelation and model identification).

### 4.3 Confidence intervals and uncertainty estimation

If we want to ask whether either of the estimated trends is statistically interesting or proceed to identify periods of significant change, we must address the issue of uncertainty in the estimated model. What uncertainty is associated with the trend estimates? One way to visualise this is through a 1 - *α*. confidence interval around the fitted trend, where *α*. is say 0.05 leading to a 95% interval. A 95% interval would be drawn at *ŷ_t_*±(*m*_1−*α*_× SE (*ŷ_t_*)), with *m*_1−*α*_ = 1.96, the 0.95 probability quantile of a standard normal distribution^1^, and SE(*ŷ_t_*) is the standard error of the estimated trend at time *x_t_*. This type of confidence interval would normally be described as *pointwise;* the coverage properties of the interval being correct for a single point on the fitted trend, but, if we were to consider additional points on the trend, the coverage would logically be lower than 1 - *α*. This is the traditional frequentist interpretation of a confidence interval. However, the GAM described here has a Bayesian interpretation (Kimeldorf and Wahba, 1970; Silverman, 1985; Wahba, 1983, 1990) and therefore the typical frequentist interpretation does not apply. Nychka (1988) investigated the properties of a confidence interval created as described above using standard errors derived from the Bayesian posterior covariance matrix for the estimated mode parameters. Such intervals have the interesting property that they have good *across-the-function* coverage when considered from a frequentist perspective. This means that, when averaged over the range of the function, the Bayesian credible intervals shown in Figure 6 have close to the expected 95% coverage. However, to achieve this some parts of the function may have more or less than 95%-coverage. Marra and Wood (2012) recently explained Nychka’s (1988) surprising results and extended them to the case of generalized models (non-Gaussian responses).

Whilst the *across-the-function* frequentist interpretation of the Bayesian credible intervals is useful, if may be important to have an interval that contains the entirety of the true function with some state probability (1 - *α*). Such an interval is known as a *simultaneous* interval. A (1 - *α*)100% simultaneous confidence interval contains *in their entirety* 1 - *α*. of all random draws from the posterior distribution of the fitted model.

Fitting a GAM involves finding estimates for coefficients of the basis functions. Together, these coefficients are distributed multivariate normal with mean vector and covariance matrix specified by the model estimates of the coefficients and their covariances respectively. Random draws from this distribution can be taken, where each random draw represents a new trend that is consistent with the fitted trend but also reflects the uncertainty in the estimated trend. This process is known as *posterior simulation*.

Figure 8 shows 20 random draws from the posterior distributions of the GAMs fitted to the Small Water and Braya-S*ϕ* data sets. In the early period of the *δ*^15^N time series many of the posterior simulations exhibit short periods of increasing and decreasing trend, balancing out to the relatively flat trend estimated by the GAM (Fig. 8a). Reflecting this uncertainty, we might expect relatively wide simultaneous intervals during this period in order to contain the vast majority of the simulated trends. Conversely, the decreasing *δ*^15^N trend starting at ~1945 is consistently reproduced in the posterior simulations, suggesting that this feature of the time series is both real and statistically significant, and that the rate of change in *δ*^15^N is relatively precisely estimated. We see a similar pattern in Figure 8b for the Braya-S*ϕ* record; the large peak in 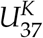 at ~250CE and the strong decline at ~1200CE are well defined in the posterior simulations, whereas most of the localised trends that are smaller magnitude changes in *y_t_* are associated with posterior simulations that are less well constrained with the ends of the record in particular showing considerable variation in the strength, timing, and even sign of simulated trends, reflecting the greater uncertainty in estimated trend during these periods. For the random draws illustrated in Figure 8, a (1 - *α*)100% simultaneous interval should contain the entire function for on average 19 of the 20 draws.

**Figure 8:**
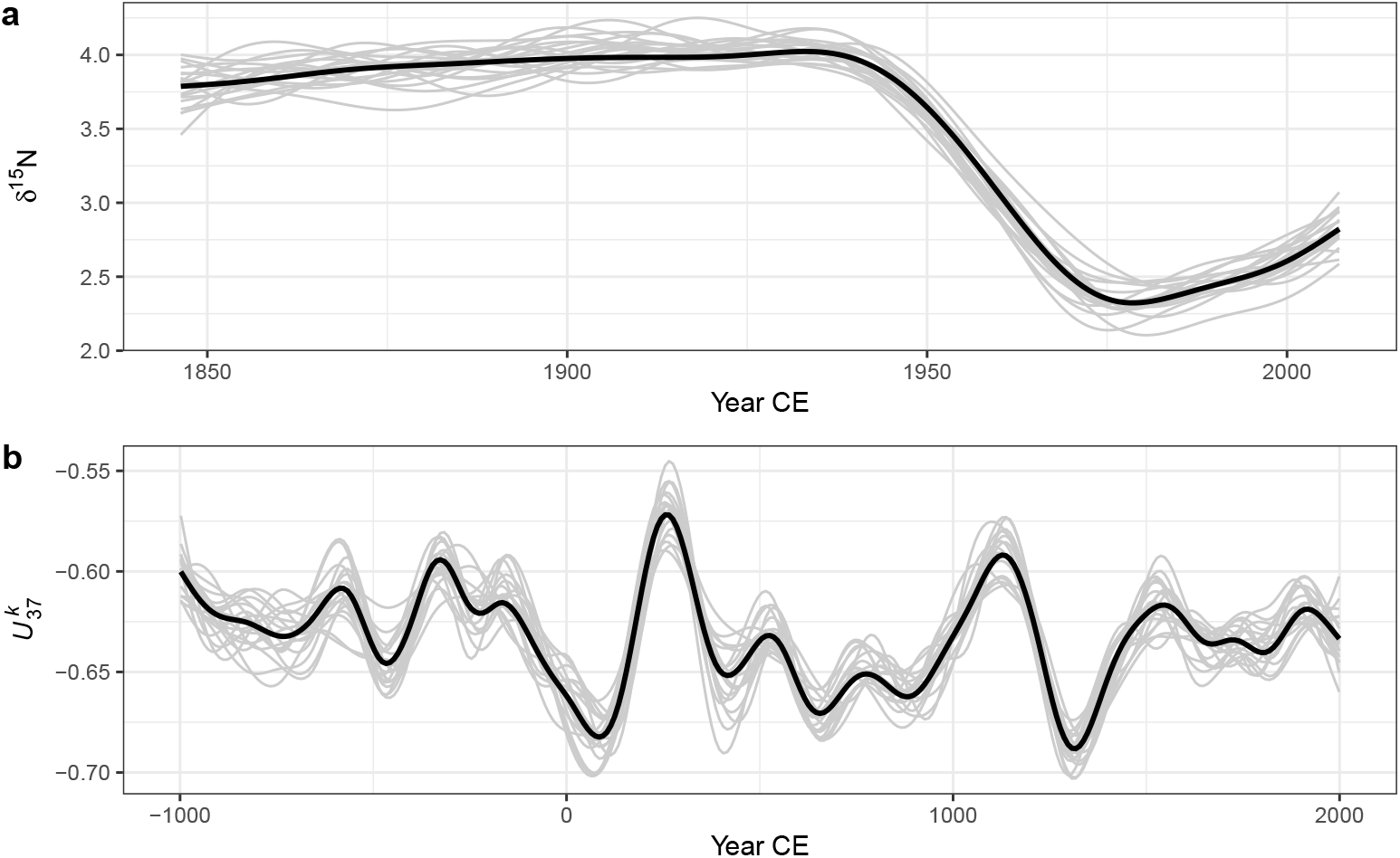
Estimated trends (thick black lines) and 20 random draws (grey lines) from the posterior distribution of the GAM fitted to the Small Water *δ*^15^N (a) and Braya-S*ϕ* 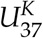 (b) time series.

There are a number of ways in which a simultaneous interval can be computed. Here I follow the simulation approach described by Ruppert et al. (2003) and present only the basic detail; a fuller description is contained in Appendix 1. The general idea is that if we want to create an interval that contains the whole of the true function with 1 - *α*. probability, we need to increase the standard Bayesian credible interval by some amount. We could simulate a large number of functions from the posterior distribution of the model and then search for the value of *m*_1−*α*_ that when multiplied by 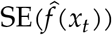 yielded an interval that contained the whole function for (1 − *α*) 100% of the functions simulated. In practice, the simulation method of Ruppert et al. (2003) does not involve a direct search, but yields the critical value *m*_1−*α*_ required.

Simultaneous intervals computed using the method described are show in Figure 9 alongside the *across-the-function* confidence intervals for the trends fitted to both example data sets. As expected, the simultaneous interval is somewhat wider than the *across-the-function* interval. The critical value *m*_1−*α*_ for the simultaneous interval of the estimated trend in *δ*^15^N is 3.07, whilst the same value for the 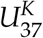 series is 3.43, leading to intervals that are approximately ±50% and ±75% wider than the equivalent across-the-function intervals.

**Figure 9:**
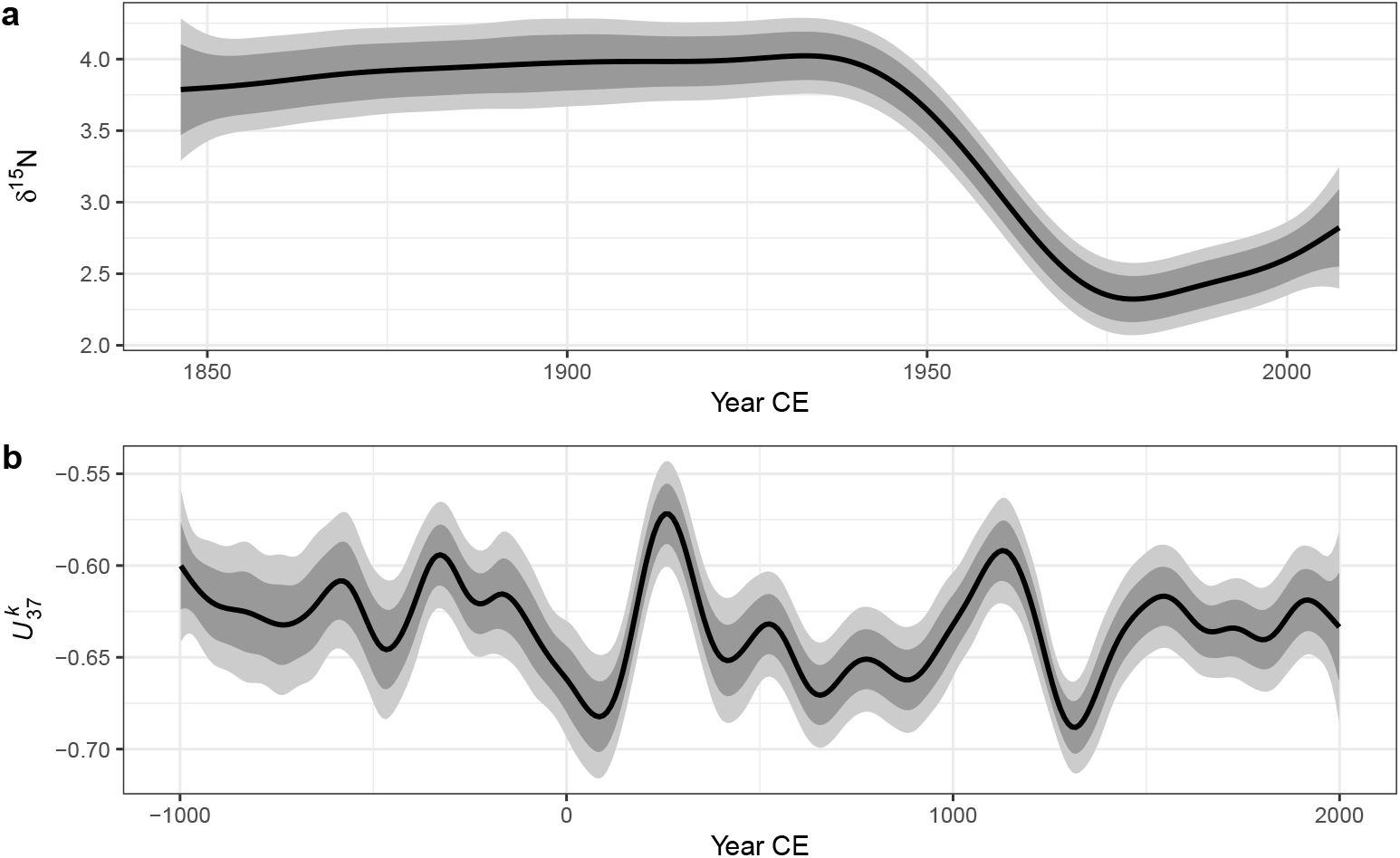
95% simultaneous confidence intervals (light grey bands) and across-the-function confidence intervals (dark grey bands) on the estimated trends (black lines) for the Small Water *δ*^15^N (a) and Braya-S*ϕ* 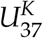 (b) time series.

### 4.4 Identifying periods change

In the simple linear trend model (1) whether the estimated trend constitutes evidence for or against a null hypothesis of no change rests on how large the estimated rate of change in *y_t_* is 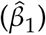 relative to its uncertainty. This is summarised in the *t* statistic. As the rate of change in *y_t_* is constant over the fitted trend — there is only a singe slope for the fitted trend 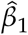 — if the *t* statistic of the test that 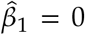 is unusually extreme this would be evidence against the null hypothesis of no change. Importantly, this applies to the whole time series as the linear model implies a constant rate of change throughout. More formally, the estimate 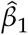 is the first derivative of the fitted trend.

In the GAM, the fitted trend need not be linear; the slope of the trend is potentially different at every point in the time series. As such we might reasonably ask *where* in the series the response *y_t_* is changing, if at all? Mirroring the linear model we can answer this question by determining whether or not the first derivative at any time point *x_t_* of the fitted trend at any time point is consistent with a null hypothesis of no change. We want to know whether or not the first derivative is indistinguishable from a value of 0 — no trend — given the uncertainty in the estimate of the derivative.

Derivatives of the fitted spline are not easily available analytically, but they can be estimated using the method of finite differences. Two values of the estimated trend, separated by a very small time-shift (Δ_*t*_), are predicted from the model; the difference between the estimated values for the two time points is an approximation of the true first derivative of the trend. As Δ_*t*_ → 0 the approximation becomes increasingly accurate. In practice, the first derivative of the fitted trend is evaluated using finite differences at a large number of points in the time series. An approximate (1 - *α*)100% pointwise confidence interval can be calculated for the derivative estimates using standard theory (i.e. ±1.96×SE(*ŷ_t_*) for a 85% interval) and the covariance matrix of the spline coefficients. A (1 - *α*)100% simultaneous interval for the derivatives can also be computed using the method described earlier. Periods of significant change are identified as those time points where the (simultaneous) confidence interval on the first derivative does not include zero.

Figure 10 shows the estimated first derivative of the fitted trend in the Small Water (10a) and Braya-S*ϕ* (10b) time series. Although the estimated trend suggests a slight increase in *δ*^15^N from the start of the record to ~1940, the estimated trend is sufficiently uncertain that the simultaneous interval on the first derivative includes 0 throughout. We can understand why this is so by looking at the posterior simulations in Figure 8a; there is considerable variation in the shape of the simulated trends throughout this period. From ~1925 the derivative of the trend becomes negative, however it is not until ~1940 that the simultaneous interval doesn’t include 0. At this point we have evidence to reject the null hypothesis of no change. This time point may be taken as the first evidence for change in *δ*^15^N in the Small Water core. The simultaneous interval on the first derivative of the trend in *δ*^15^N is bounded away from 0 between ~1940 and ~1975, covering the major decline in values evident in the observations. The simultaneous interval includes 0 from ~1975 onward, suggesting that, whilst quite pronounced, the recent increase in *δ*^15^N is not statistically significant. To determine whether or not the recent increase is real, we would require considerably more samples with which to (hopefully) more-precisely estimate the trend during this period. Alternatively, we might just have to wait until sufficient additional sedimentation has occurred to warrant recoring Small Water and reestimating the trend in *δ*^15^N.

The estimated trend at Braya-S*ϕ* exhibited a number of oscillations in 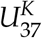. As we saw previously in Figures 8b and 9b, many of these are subject to significant uncertainty and it is important therefore to discern which, if any, of the oscillations in the response can be identified given the model uncertainty. In Figure 10b only two features of the estimated trend are considered significant based on the derivatives of the smooth; one centred on ~250CE and a second at ~1150CE. In both these periods, the simultaneous interval for the first derivative of the trend does not include zero. In the first case we detect the large peak and subsequent decline in 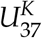 at ~250CE, whilst at ~1150CE the large trough is identified, but not the increasing trend immediately prior to this excursion to lower 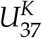. Recall that these intervals are simultaneous in nature, strongly guarding against false positives, and as such we can be confident in the estimation of these two features, whilst care must be taken to not over-interpret the remaining variations in the estimated trend.

**Figure 10:**
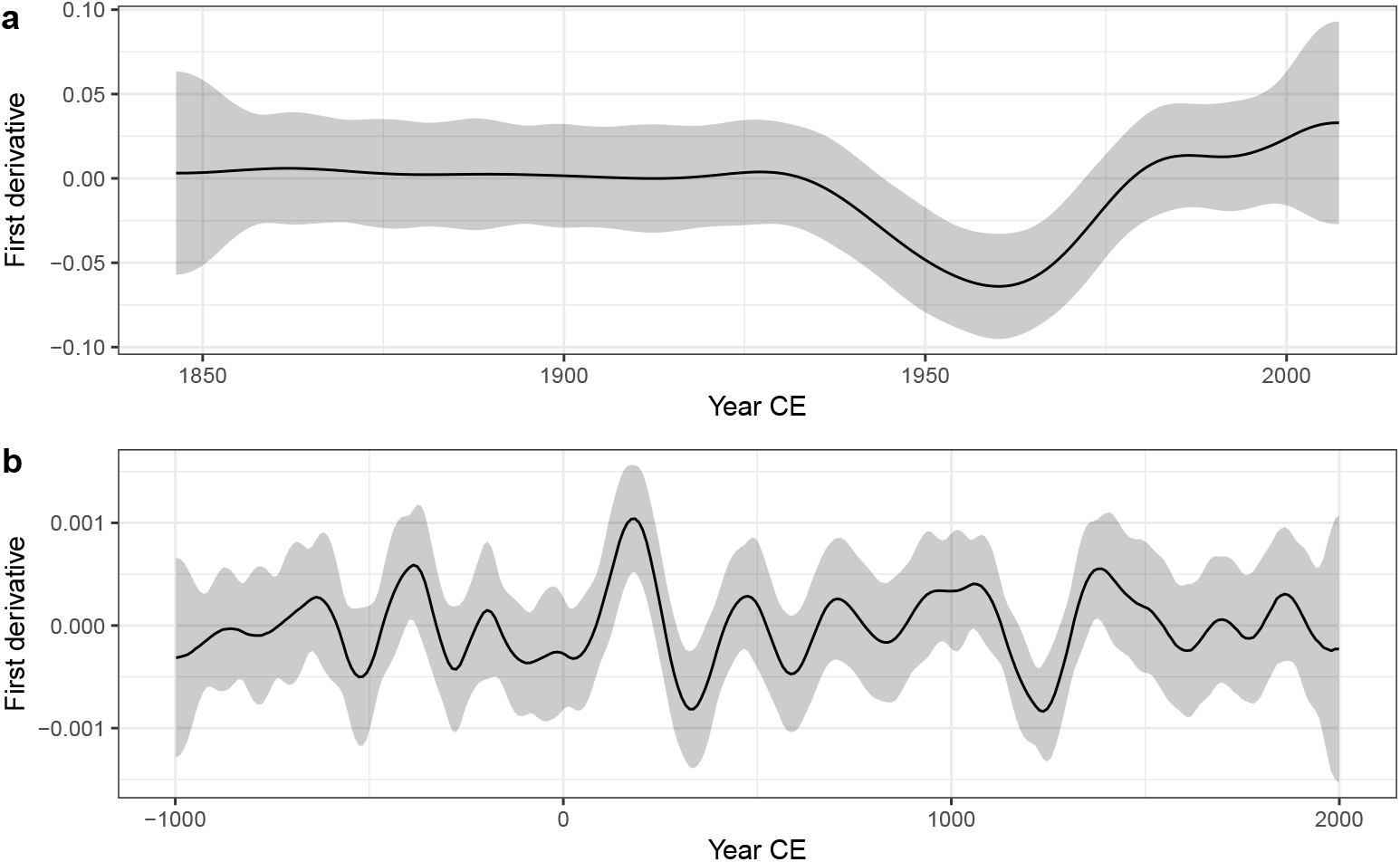
Estimated first derivatives (black lines) and 95% simultaneous confidence intervals of the GAM trends fitted to the Small Water *δ*^15^N (a) and Braya-S*ϕ* 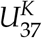 (b) time series. Where the simultaneous interval does not include 0, the models detect significant temporal change in the response.

### 4.5 Residual autocorrelation and model identification

The GAM fitted to the *δ*^15^N time series contained a CAR(1) process to model residual temporal autocorrelation in the residuals. The estimated magnitude of the autocorrelation is given by the parameter *ϕ*. The estimated value of *ϕ* for the *δ*^15^N series is 0.6 with 95% confidence interval 0.28–0.85, indicating moderate to strong residual autocorrelation about the fitted trend. The correlation function is an exponentially decreasing function of temporal separation (Δ_*t*_), and whilst observations that are a few years apart are quite strongly dependent on one another, this dependence drops off rapidly as Δ_*t*_ increases and is effectively zero when samples are separated by a decade or more (Figure 11).

**Figure 11:**
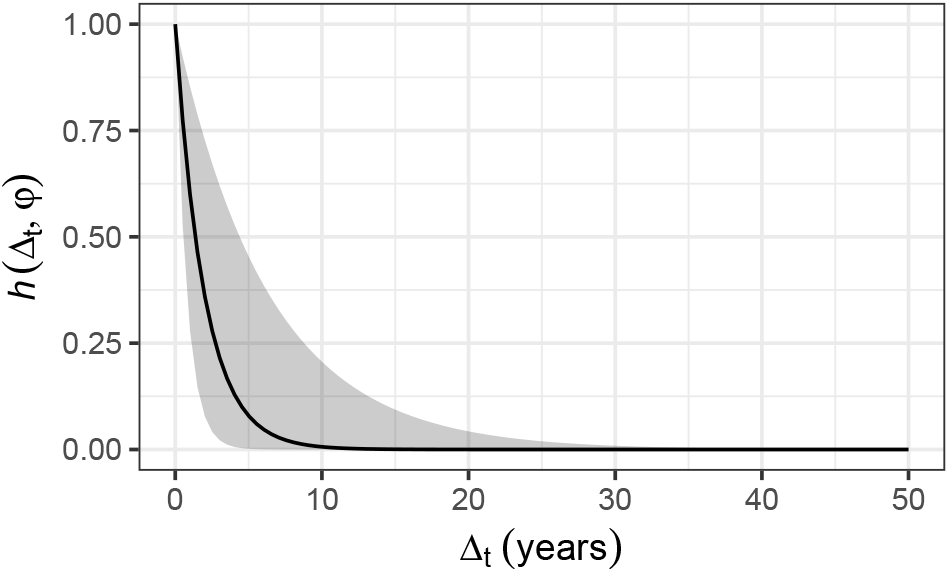
Estimated CAR(1) process from the GAM fitted to the Small Water *δ*^15^N time series. *h*(Δ_*t*_, *ϕ*) is the correlation between residuals separated by *Δ_t_* years, where 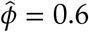 The shaded band is a 95% pointwise confidence interval on the estimated correlation *h*.

Failure to account for the dependencies in the *δ*^15^N time series could lead to the estimation of a more wiggly trend than the one shown in Figure 6a which would negatively impact the confidence placed on the inferences we might draw from the fitted model. Importantly, failing to account for the strong dependency in the residuals would lead to smaller uncertainties in the estimated spline coefficients, which would propagate through to narrower confidence intervals on the fitted trend and on the first derivatives, and ultimately to the identification of significant periods of change. The end result would be a tendency toward anti-conservative identification of periods of change; the coverage probability would be lower than the anticipated 1 – *α*, leading to a greater chance of false positive results.

Problems estimating the GAM plus CAR(1) model were encountered when this was fitted to the 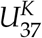 time series; including both a smooth trend in the mean 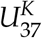 and a CAR(1) process in the residuals lead to an unidentifiable model. What makes a model with a spline-based trend and an autocorrelation process like the CAR(1) potentially unidentifiable?

Consider again the basic GAM for a smooth trend, (3). In that equation the correlation matrix Λ was omitted for the sake of simplicity. As I did in (6), I reintroduce it and restate the distributional assumptions of this model

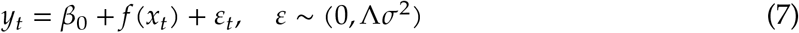

In the basic GAM, Λ ≡ **I** is an identity matrix, a matrix with 1s on the diagonal and 0s elsewhere

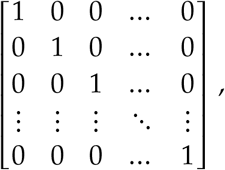

which is where the independence assumption of the model comes from; a model residual is perfectly correlated with itself (the 1s on the diagonal), but uncorrelated with any other residual (the off-diagonal 0s). In the GAM plus CAR(1) model, an alternative correlation function for Λ was used — the CAR(1) with correlation parameter *ϕ*. Fahrmeir and Kneib (2008) show that where the stochastic structure of *f* and Λ approach one another, i.e. where we have a potentially wiggly trend or strong autocorrelation as *ϕ* → 1, the two processes can quickly become unidentifiable (see also Fahrmeir et al., 2013). By unidentifiable, we mean that it becomes increasingly difficult to distinguish between a wiggly trend or strong autocorrelation because these two processes are very similar to one another in appearance. This leads to model estimation problems of the sort encountered with fitting the GAM plus CAR(1) model to the Braya-s*ϕ* 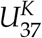 series.

Why might this be so? Autocorrelation is the tendency for a large (small) value of *y_t_* at time *x_t_* to be followed by a likewise large (small) value at time *x*_*t*+1_. This leads to runs of values that are consistently greater (less) than the overall mean. Short runs would indicate weaker autocorrelation whilst longer runs are associated with stronger autocorrelation, and long runs of values greater (less) than the mean would be evident as non-linear trends in the time series. As a result, a wiggly trend and an autocorrelation function with large *ϕ* are two ways to describe the same pattern of values in a time series, and, without any further information to constrain either of these, the model is unable to distinguish both components uniquely.

Situations where it may be possible to uniquely identify separate wiggly trends and autocorrelation are exemplified by the Small Water *δ*^15^N time series. The non-linear trend and the autocorrelation operate at very different scales; the trend represents decadal-scale variation in mean *δ*^15^N, whilst the CAR(1) process represents the much smaller-scale tendency for values of the response to be followed in time by similar values. That such a pattern is observed in the Small Water core is the result of the high resolution of the sampling in time relative to the long-term trend. In contrast, the Braya-S*ϕ* record is sampled at far lower resolution relative to the fluctuations in the mean response, and consequently the data do not contain sufficient information to separate trend and autocorrelation.

### 4.6 Gaussian process smooths

In the world of machine learning, Gaussian processes (Golding and Purse, 2016; Rasmussen and Williams, 2006) are a widely-used method for fitting smooth non-parametric regression models. A Gaussian process is a distribution over all possible smooth functions *f*(*x*). In the field of spatial statistics, Gaussian processes are known by the name *kriging*.

With a Gaussian process we are interested in fitting a smooth temporal trend by modelling the way the correlation between pairs of observations varies as a function of the distance, *h*, in time that separates the observations. The correlation between pairs of observations decreases with increasing separation, which is modelled using a correlation function, *c*(*h*).

Several functions can be used to represent *c*(*h*). Two common ones are the power exponential function and the Matérn family of correlation functions. The power exponential function at separation distance *h* is

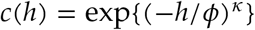

where 0 < *κ* ≤ 2. The Matérn correlation function is actually a family of functions with closed-forms only available for a subset of the family, distinguished by *κ*. When *κ* = 1.5, the Matérn correlation function is

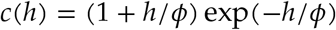

whilst for *κ* = 2.5 it is

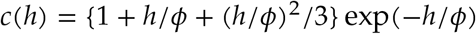

and for *κ* = 3.5

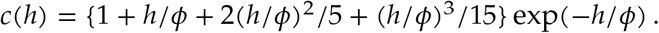

In all cases, *ϕ* is the effective range parameter, which sets the distance beyond which the correlation function is effectively zero.

Figure 12 shows examples of two different correlation functions; the *power exponential* (Figure 12a), and the Matérn (Figure 12b) correlation functions. These functions are smooth and monotonic-decreasing, meaning that the value of the correlation function decreases with increasing separation (*h*). When *h* = 0, the correlation is equal to 1 (*c*(0) = 1); two samples taken at exactly the same time point are perfectly correlated. As *h* → ∞, the correlation tends to zero (*c*(*h*) → 0); two samples separated by a large amount of time tend to be uncorrelated. Often we are interested in learning how large the separation in time needs to be before, on average, a pair of observations is effectively uncorrelated (i.e. where *c*(*h*) is sufficiently close to zero).

**Figure 12:**
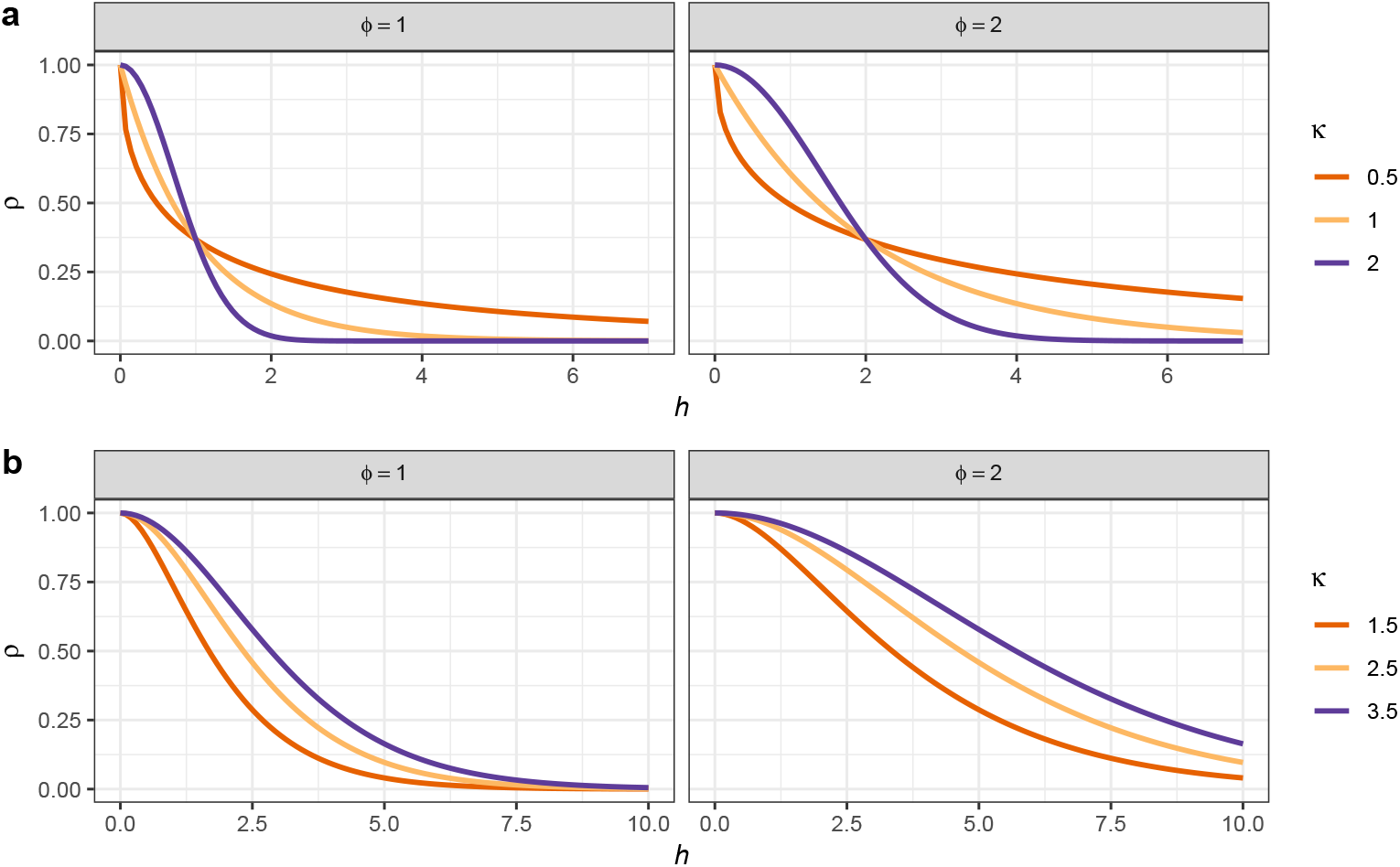
Power exponential (a) and Matérn (b) correlation functions for observation separation distance *h*. Two values of the effective range parameter (*ϕ*)) are shown for each function. For the power exponential function, *κ* is the power in the power exponential function. For the Matérn correlation function, *κ* distinguishes the member of the Matérn family.

Gaussian processes and GAMs share many similarities and we can fit a Gaussian process using the techniques already described above for splines (Handcock et al., 1994; Kammann and Wand, 2003). It can be shown (e.g. Fahrmeir et al., 2013) that the Gaussian process model has the same penalised likelihood form as the GAM that we discussed earlier; the observations are the knots of the smoother and each has a basis function in the form of a correlation function. The equivalence is only true if the basis functions do not depend on any other parameters of the model, which is only achievable if the value of *ϕ* is fixed and known (Fahrmeir et al., 2013). In general, however, we would like to estimate *ϕ* as part of model fitting. To achieve this we can maximise the profile likelihood or score statistic of the model over a range of values of *ϕ* (Wood, 2017, 362–363). This involves proposing a value of *ϕ* for the effective range of the correlation function and then estimating the resulting GAM by minimising the penalised log-likehood conditional upon this value of *ϕ* and repeating for a range of values for *ϕ*. The model, and its corresponding value of *ϕ*, with lowest penalised log-likelihood or score statistic is then retained as the estimated GAM. Figure 13a shows the REML score for models estimated using a Gaussian process smooth with a Matérn correlation function (*κ* = 1.5) for a sequence of values of *ϕ* between 15 and 1000 years. There is a clear minimum around 40 years separation, with the minimum REML score being observed at *ϕ* = 41.81). Also shown are the REML scores for models using the power exponential function (*κ* = 1) with the minimum score observed at a somewhat higher effective range of *ϕ* = 71.06.

**Figure 13:**
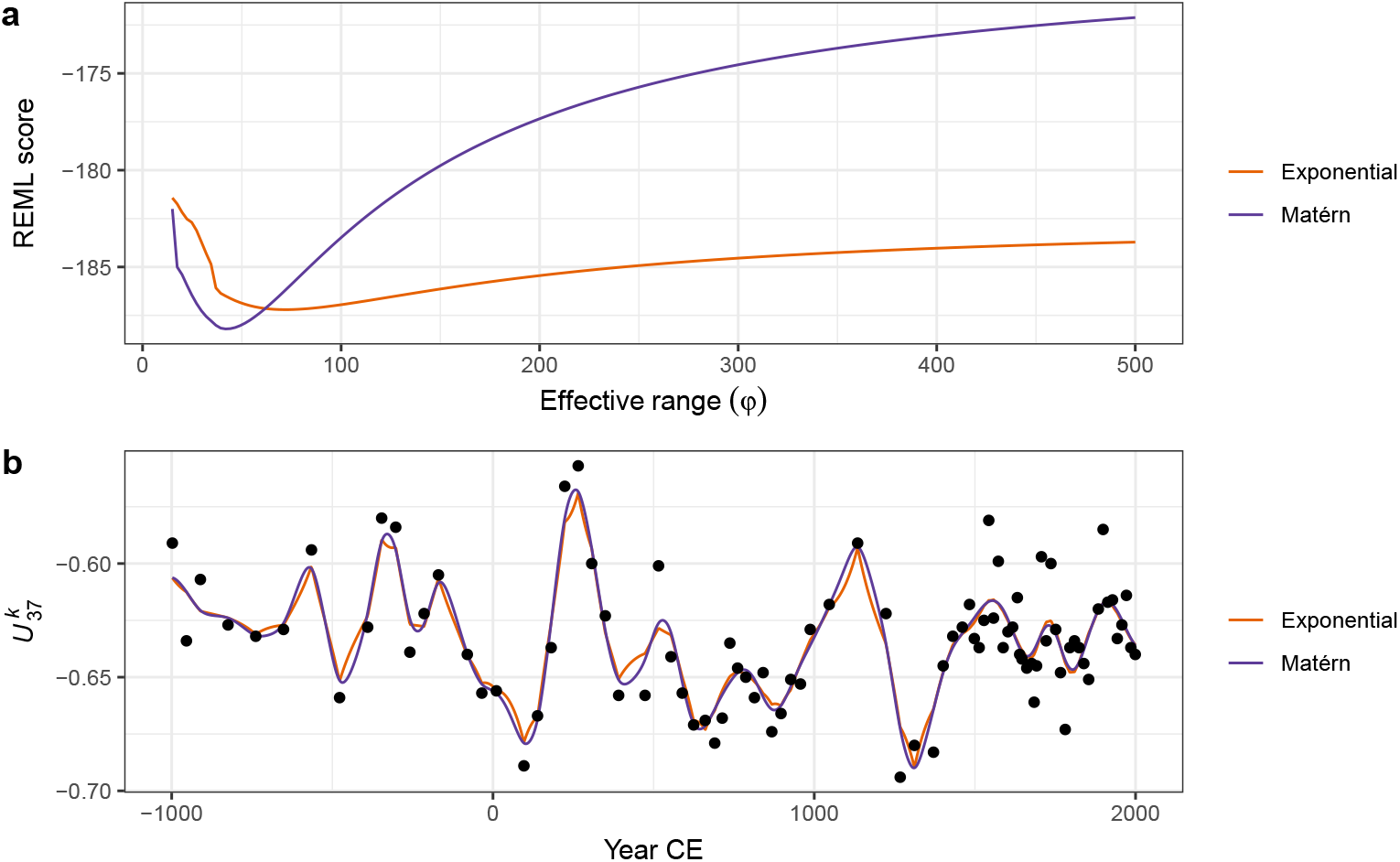
Gaussian process smooths fitted to the 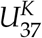 time series. REML score traces for GAMs fitted using power exponential (*κ* = 1) or Matérn (*κ* = 1.5) correlation functions as a function of the effective range parameter (*ϕ*) are shown (a). The optimal model for each function is that with the lowest REML score. b) shows the resulting trends estimated using the respective correlation function with the value of *ϕ* set to the optimal value.

Figure 13b shows the estimated trends for the 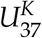 time series using Gaussian process smooths with exponential and Matérn correlations functions, both using *ϕ* values at their respective optimal value as assessed using the REML score. The estimated trends are very similar to one another, although there is a noticeable difference in behaviour, with the power exponential (*κ* = 1) version being noticeably less-smooth than the Matérn version. This difference is attributable to the shapes of the respective correlation functions; the Matérn approaches a correlation of 1 smoothly as *h* approaches 0, whilst the power exponential with *κ* = 1 approaches a correlation of 1 increasingly quickly with decreasing h. The power exponential with *κ* = 2, like the Matérn, approaches *ϕ* = 1 smoothly, and consequently the trend estimated using this correlation function is qualitatively similar to that estimated using the Matérn correlation function.

### 4.7 Adaptive smoothing

Each of the spline types that I have discussed so far shares a common feature; the degree of wiggliness over the time series is fixed due to the use of a single smoothness parameter, *λ*. The definition of wiggliness, as the integrated squared second derivative of the spline, ensures that the fitted smoother does not jump about wildly. This assumes that the data themselves are well described by a smoothly varying trend. If we anticipate abrupt change or step-like responses to environmental forcing this underlying assumption of the GAM would suggest that the method is ill-suited to modelling palaeo time series in which such features are evident or expected.

While there is not much we can do within the GAM framework to model a series that contains both smooth trends and step-like responses, an adaptive smoother can help address problems where the time series consists of periods of rapid change in the mean combined with periods of complacency or relatively little change. As suggested by their name, adaptive smoothers can adjust to changes in the wiggliness of the time series. This adaptive behaviour is achieved by making the smoothness parameter *λ* itself depend smoothly on *x_t_* (Ruppert et al., 2003, 17; Wood, 2017, 5.3.5); in other words, the adaptive smoother allows the wiggliness of the estimated trend to vary smoothly over time. Whilst this allows the estimated trend to adapt to periods of rapid change in the response, adaptive smoothers make significant demands on the data (Wood, 2017, 5.3.5); if we used *m* smoothness penalties to allow the wiggliness to vary over a time series, it would be like estimating *m* separate smooths from chunks of the original series each of length *n/m*. In a practical sense, this limits the use of adaptive splines in palaeoecology to proxies that are readily enumerated, such as the biogeochemical proxies used in the two example data sets.

Figure 14 compares trends for the Braya-S*ϕ* time series estimated using GAMs with the three main types of spline discussed; i) TPRS, ii) Gaussian process smooths, and iii) an adaptive smoother using 45 basis functions and 5 smoothing parameters. There is a clear difference in the behaviour of the adaptive and non-adaptive smoothers for the first 1000 years of the record, with the adaptive smooth exhibiting much less variation compared with either the TPRS or Gaussian process splines. Over the remaining two thirds of the series, there is much closer agreement in the three smooths.

**Figure 14:**
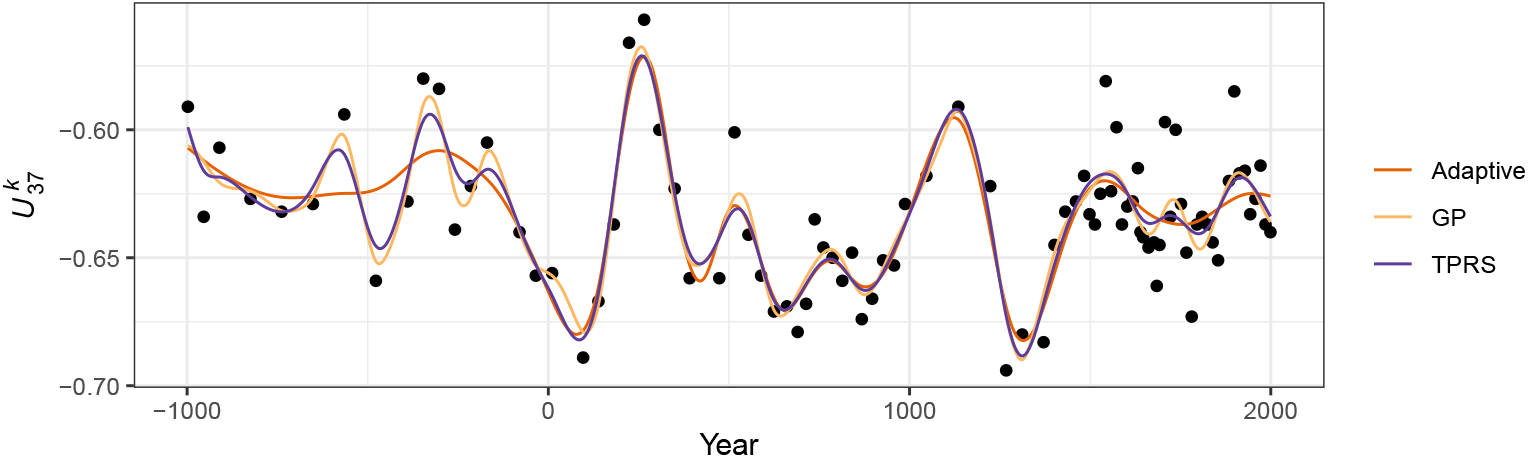
Comparison of trends estimated using i) adaptive smooth, ii) Gaussian process, and iii) thin plate regression spline bases for the 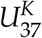 time series.

The behaviour of the TPRS and Gaussian process splines for these data is the result of requiring a large amount of wiggliness (a small *λ*) to adapt to the large oscillations in 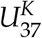 present around year 250CE and again at ~900–1500CE. This large degree of wiggliness allows the splines to potentially over-fit individual data points much earlier in the record. Because the adaptive smoother, in contrast, can adapt to these periods of rapid change in the response it is much less susceptible to this “chasing” behaviour — we don’t need to waste effective degrees of freedom in periods with little or no change just to be able to fit the data well when there is a lot of change.

This potential for over-fitting in such situations is undesirable, yet if we recall Figure 10 and the discussion around the use of the first derivative to identify periods of significant change, we would not interpret the oscillations in the early part of the 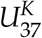 record as being statistically significant. Owing to the paucity of data in this part of the series the trends fitted using the non-adaptive smoothers are subject to such a large degree of uncertainty that the alternative of no trend through the first 1000 years of the record is also a plausible explanation of the data. The trend estimated using the adaptive smooth reflects this. Therefore, should we conclude that there is no trend in 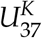 and thence climate in this period? I believe that to be too-strong a statement; those oscillations in 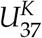 may be real responses to climate forcing but we may simply lack the statistical power to distinguish them from the null hypothesis of no trend through this period. The adaptive smoother is only adjusting to the data available to it; just because it does not detect a trend during this period does not lend itself to an interpretation of stable climate forcing or complacency in the lake’s response to forcing (although that is a justifiable interpretation of the result). If there were particular interest in the climate of this particular period we might take from the Braya-S*ϕ* record that there is potential early variation due to climate forcing, but that additional data from this or other sites is required before any definitive conclusion can be drawn.

### 4.8 Accounting for age model uncertainty

Thus far, the trend models that I have described and illustrated assumed that the time co-variate (*x_t_*) was fixed and known. In both examples, and generally for most palaeoecological records, this assumption is violated. Unless the record is annually laminated, assigning an age to a sediment interval requires the development of an age model from observations of the relationship between depth down the sediment core and estimates of the age of the sample arrived at using any of a number of techniques, for example ^210^Pb or ^14^C radiometric dating. This age-depth relationship is itself uncertain, usually being derived from a mathematical or statistical model applied to point age estimates (e.g. Blaauw and Heegaard, 2012). Incorporating this additional component of uncertainty complicates the estimation of statistical models from palaeoenvironmental data. In this section I illustrate a simulation based approach to quantify and account for age-model uncertainty as part of the trend estimation using a GAM (see Anchukaitis and Tierney (2013) for a similar, non-GAM related idea).

Figure 15a shows the estimated dates (in Years CE) for 12 levels in the Small Water core dated using ^210^Pb. The vertical bars show the estimated age uncertainty of each level. The solid line through the data points is an additive model fitted to the observations, with prior weights given by the estimated age uncertainties. The fitted age-depth model is constrained to be monotonically decreasing with increasing depth, following the method of (Pya and Wood, 2015) using the *scam* package (Pya, 2017). Also shown are 25 simulations from the posterior distribution of the monotonically-constrained GAM. Each simulation from the posterior distribution of the age-model is itself a potential age-depth model, which can be used to assign dates to the Small Water core. The trend model in (4) can be fitted to the *δ*^15^N data using these new dates as *x_t_*, and the whole process repeated for a large number of simulations from the age model.

**Figure 15:**
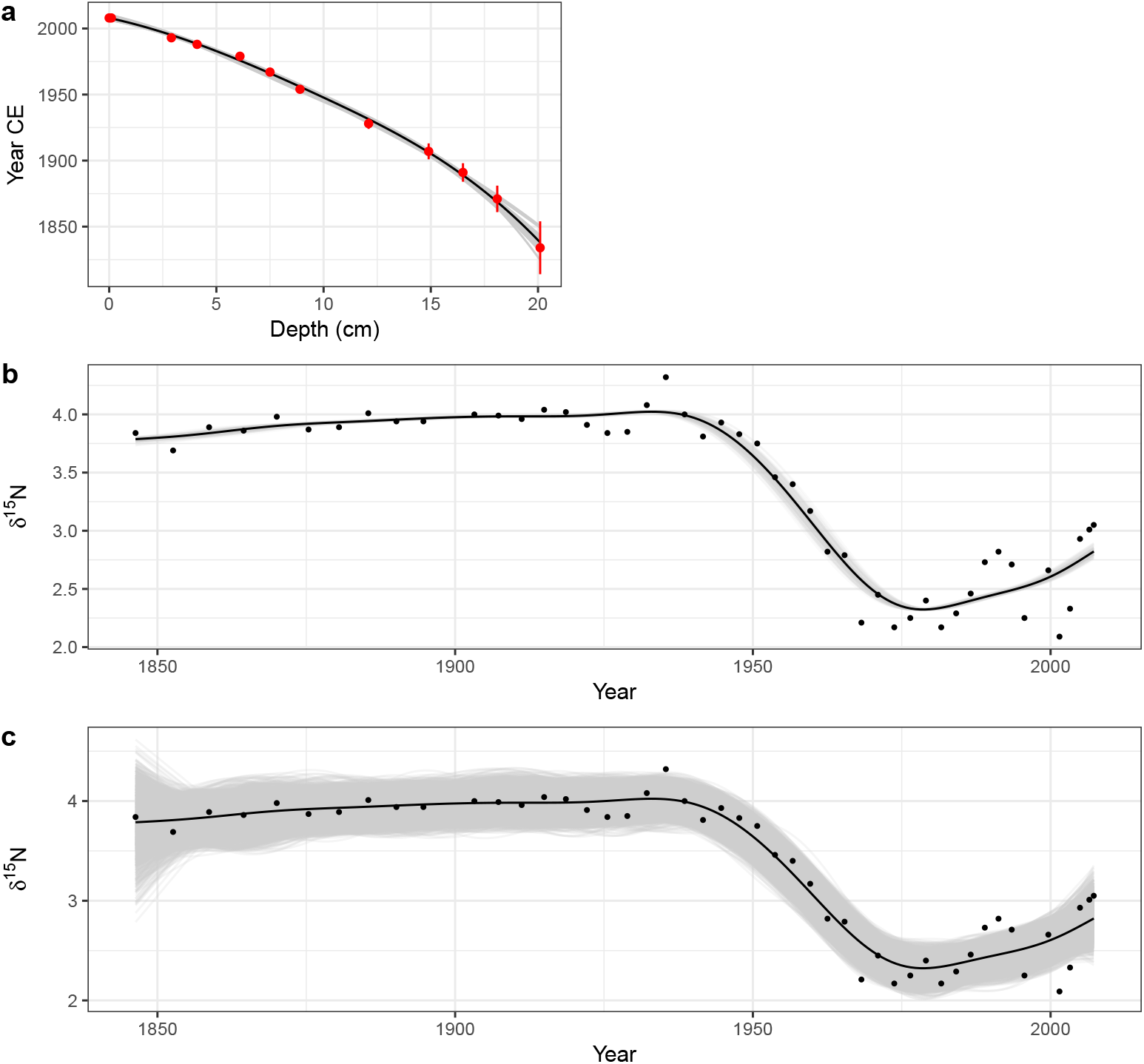
Accounting for uncertainty in age estimates whilst fitting a smooth trend to the Small Water *δ*^15^N time series. (a) Estimated age model using a monotonically-constrained spline fitted to ^210^Pb inferred ages for selected depths in the sediment core (red points). The uncertainty in the ^210^Pb inferred age is show by the red vertical bars. The fitted age model is illustrated by the solid black line. The faint grey lines are 25 random draws from the posterior distribution of the monotonically constrained GAM. The effect of age uncertainty on trend estimation is shown in b); for 100 simulations from the posterior distribution of the age model in a) a trend was estimated using a GAM with a thin plate regression spline basis and a CAR(1) process in the residuals. These trends are shown as grey lines. The combined effect of age model and fitted GAM uncertainty on the trends for the *δ*^15^N time series is shown in c). The grey lines in c) are based on 50 random draws from the model posterior distribution for each of the 100 trends shown in b). For both b) and c) the black line shows the trend estimated assuming the ages of each sediment sample are known and fixed.

Figure 15b shows the trend in *δ*^15^N for the observed age-depth model, plus trends estimated via the same model using 100 draws from the posterior distribution of the age model. In this case, the age-depth model is relatively simple with little variation in the posterior draws, resulting in trends that match closely that obtained from the estimated age-depth relationship. Even so, this additional uncertainty suggests that the timing of the decline in *δ*^15^N covers the interval ~1935–1945.

The uncertainty in the trend estimates illustrated in Figure 15b only reflects the variation in trends fitted to the uncertain dates of the sediment samples. To fully visualise the uncertainty in the trend estimates, incorporating both age model uncertainty *and* uncertainty in the estimated model coefficients themselves, 50 simulations from the posterior distribution of each of the 100 estimated trends shown in Figure 15b were performed, resulting in 5,000 trend estimates for the *δ*^15^N series. These are shown in Figure 15c, where the two obvious changes over the same simulations without accounting for uncertainty in *x_t_* (Figure 8a) are that the uncertainty band traced out by the simulations is approximately 50% wider and, not surprisingly, the uncertainty in the estimated trend is most pronounced in the least accurately-dated section of the core. Despite this additional uncertainty however, the main result holds; a marked decline of ~1.5‰ that occurred between approximately 1930 and 1945, with mild evidence of a small increase in *δ*^15^N post 2000 CE.

### 4.9 Multivariate data

A large proportion of the palaeoenvironmental data generated today is multivariate in nature and yet the two examples used to illustrate GAMs were univariate. Can the approach described here be used for multivariate data? Yes, and no. With one main exception it is not possible to directly apply the GAM methodology described here to multivariate abundance data, where the aim is to model all species at once. The *mgcv* software, for example, is not able to estimate the penalized GAM for multiple non-Gaussian responses. The exception is for a small number of correlated Gaussian responses; these could be modelled as being distributed multivariate normal conditional upon the covariates. Such a model would estimate the expected values of each response and the correlations between them. For example, we could jointly model *δ*^15^N and *δ*^13^C series using this approach.

Formal multivariate versions of GLM or GAMs are currently an important area of research within ecology (see Warton et al. (2015) for a recent review), where they go by the name joint species distribution models (JSDMs). Whilst undoubtedly powerful, our knowledge regarding JSDMs and their availability in software are still in their relative infancy and they require considerable expertise to implement. As such, JSDMs are currently beyond the reach of most palaeoecologists. Despite this, we should be watching JSDM research as developments are ongoing and a degree of method maturation occurring.

One currently available avenue for fitting a multivariate GAM is via regularized sandwich estimators and GLMs (Warton, 2011), which involves fitting separate GLMs (or GAMs) to each response variable and subsequently using resampling-based hypothesis tests to determine which covariates are related to variation at the community level and for individual taxa (Wang et al., 2012; Warton, 2011; Warton et al., 2012). The *mvabund* package (Wang et al., 2012) implements this approach within R and can use *mgcv* to fit GAMs to each species.

A pragmatic although inelegant approach that has been used to estimate trends in multivariate palaeoecological data is to first summarise the response data using an unconstrained ordination via a PCA, CA, or principal curve and then fit separate GAM models to the site (sample) scores of the first few ordination axes or principal curve (Beck et al., 2018; Bennion et al., 2015).

Whilst this two-step approach is relatively easy to implement and builds on approaches that palaeoecologists already use to summarise multivariate stratigraphic data, it is best thought of as modelling changes in abundance or relative composition at the community level. It is less well suited to unpicking taxon-specific trends however, because the ordination step combines individual species information into latent variables (axes) that are linear combinations of *all* species and it is these latent variables that are then modelled using GAM.

## 5 Conclusions

Formal statistical estimation of trends in palaeoenvironmental data has been hampered by the nature of the data that comprise the time series; the uneven spacing of samples in time makes it, if not impossible, difficult to fit classical statistical time series models like ARIMA. This has led palaeoecologists and palaeolimnologists to either ignore statistical estimation of trends or fall back on basic statistical methods such as linear parametric and non-parametric correlations or simple linear regression models, where the assumptions of the method are often grossly violated by the dependencies inherent to time series data. GAMs, whilst similar to the popular Loess smoother, provide a superior alternative approach to trend estimation in palaeoenvironmental time series. GAMs can estimate non-linear trends, provide estimates of the magnitude of change as well as allow the identification of periods of change, can account for the lack of independence (either via autocorrelation processes or via the fitting of a wiggly trend), and provide a formal framework for statistical inference on each of these features.

In presenting the GAM with specific palaeoenvironmental examples and addressing the issues that arise in palaeoenvironmental time series, it is hoped that palaeoecologists and palaeolimnologists will be motivated to give greater consideration to the estimation of trends and the identification of change in stratigraphic time series.

## Acknowledgements

The ideas expressed in this paper are the result of many fruitful conversations with colleagues past and present at the Environmental Change Research Centre, UCL, and the University of Regina. In particular I am indebted to Helen Bennion, Rick Battarbee, and Peter Leavitt for their collaborations on projects over many years, and to David Miller, Eric Pedersen, and Noam Ross, my GAM workshop partners in crime. Without Simon Wood’s *mgcv* software and his research on GAMs, the application of these models to palaeo time series would not be as straight forward. This work was supported by a Natural Sciences and Engineering Council of Canada (NSERC) Discovery Grant to the author (RGPIN-2014-04032).

## Appendix 1 Simultaneous intervals

We proceed by considering a simultaneous confidence interval for a function *f*(*x*) at a set of *M* locations in *x*; we’ll refer to these locations, following the notation of Ruppert et al. (2003) by

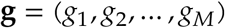

The true function over **g, f_g_**, is defined as the vector of evaluations of *f* at each of the *M* locations

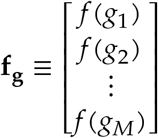

and the corresponding estimate of the true function given by the fitted GAM denoted by 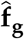. The difference between the true function and our unbiased estimator is given by

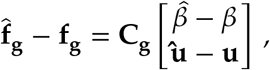

where **C_g_** is a matrix formed by the evaluation of the basis functions at locations **g**, and the part in square brackets is the bias in the estimated model coefficients, which we assume to be mean **0** and distributed, approximately, multivariate normal with mean vector 0 and covariance matrix **V_b_**

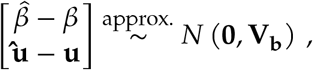

where **V_b_** is the Bayesian covariance matrix of the GAM coefficients.

Now, the (1 - *α*)100% simultaneous confidence interval is

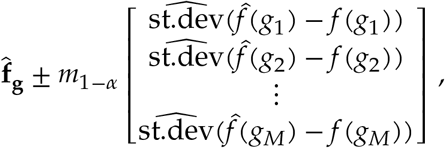

where *m*_1-*α*_ is the 1 - *α*. quantile of the random variable

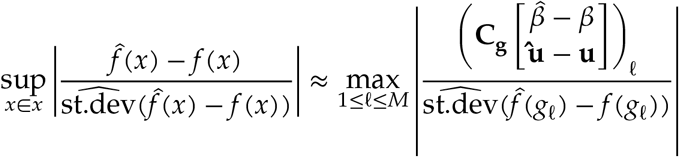

The sup refers to the *supremum* or the *least upper bound;* this is the least value of 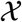, the set of all values of which we observed subset *x*, that is *greater* than all of the values in the subset. Often this is the maximum value of the subset. This is what is indicated by the right-hand side of the equation; we want the maximum (absolute) value of the ratio over all values in **g**.

The fractions in both sides of the equation correspond to the standardized deviation between the true function and the model estimate, and we consider the *maximum absolute* standardized deviation. We don’t usually know the distribution of the maximum absolute standardized deviation but we need this to access its quantiles. However, we can closely approximate the distribution via simulation. The difference here is that rather than simulating from the posterior of the model as we did earlier see section *Confidence intervals*, this time we simulate from the multivariate normal distribution with mean vector 0 and covariance matrix **V_b_**. For each simulation we find the maximum absolute standardized deviation of the fitted function from the true function over the grid of *x* values we are considering. Then we collect all these maxima, sort them and either take the 1 - *α*. probability quantile of the maxima, or the maximum with rank ⌈(1 - *α*)/*N*⌉.

## Footnotes

1 The 0.95 probability quantile of the *t* distribution may be used instead, which will account for estimation of *σ*, the variance of the data. However, given the number of observations, and hence residual degrees of freedom, needed to motivate fitting GAMs, differences between intervals computed using extreme quantiles of the standard normal or the *t* distribution will be tiny.

